# Imputation of single-cell gene expression with an autoencoder neural network

**DOI:** 10.1101/504977

**Authors:** Md. Bahadur Badsha, Rui Li, Boxiang Liu, Yang I. Li, Min Xian, Nicholas E. Banovich, Audrey Qiuyan Fu

## Abstract

**Background:** Single-cell RNA-sequencing (scRNA-seq) is a rapidly evolving technology that enables measurement of gene expression levels at an unprecedented resolution. Despite the explosive growth in the number of cells that can be assayed by a single experiment, scRNA-seq still has several limitations, including high rates of dropouts, which result in a large number of genes having zero read count in the scRNA-seq data, and complicate downstream analyses.

**Methods:** To overcome this problem, we treat zeros as missing values and develop nonparametric deep learning methods for imputation. Specifically, our LATE (Learning with AuToEncoder) method trains an autoencoder with random initial values of the parameters, whereas our TRANSLATE (TRANSfer learning with LATE) method further allows for the use of a reference gene expression data set to provide LATE with an initial set of parameter estimates.

**Results:** On both simulated and real data, LATE and TRANSLATE outperform existing scRNA-seq imputation methods, achieving lower mean squared error in most cases, recovering nonlinear gene-gene relationships, and better separating cell types. They are also highly scalable and can efficiently process over 1 million cells in just a few hours on a GPU.

**Conclusions:** We demonstrate that our nonparametric approach to imputation based on autoencoders is powerful and highly efficient.

## INTRODUCTION

Due to dropout and other technical limitations in single-cell sequencing technologies, single-cell RNA-seq (scRNA-seq) data typically contain many zero expression values. This is particularly true for droplet-based scRNA-seq technologies (such as 10x Genomics), which are the most commonly used in the field (see review in Kolodziejczyk, et al. [1]). This often results in a read count of zero for over 80% (sometimes over 90%) of all the measurements across genes and across cells [2]. The high rate of zeroes leads to difficulties in downstream analyses: for example, it may obscure gene-gene relationships, or blur differences in subpopulations of cells (i.e., cell types). Although some of the zeroes represent no expression, most are due to failures to capture the transcript and do not indicate the true expression level. Recovering true expression levels behind the zeros in scRNA-seq data is therefore of great interest [3-7]. In this work, we treat the zeros as missing values (see comments on this assumption in Discussion) and aim to use a computational approach to impute these missing values using information available from nonzero values, exploiting the dependence in gene expression levels across genes and across cells. The imputed data will have better quality than the original data, and may be used for diverse downstream analyses. For example, using the imputed scRNA-seq data, one may perform clustering analysis to identify cell types, and differential expression analysis to identify genes that are expressed differently in different cell types, study relationships between certain genes, or construct a co-expression network of genes in single cells [8, 9].

Here, we developed imputation methods based on autoencoders for scRNA-seq data. Autoencoders [10] are a type of architecture commonly used in deep learning and enable reconstruction of the input through dimension reduction. Our LATE (Learning with AuToEncoder) method trains an autoencoder *de novo* on the highly sparse scRNA-seq data, with the initial values of the parameters randomly generated. Our TRANSLATE (TRANSfer learning with LATE) method builds on LATE and further incorporates a reference gene expression data set (e.g., bulk gene expression, a larger scRNA-seq data set, data from a complementary scRNA-seq technology, or scRNA-seq data of similar cells types collected elsewhere) through transfer learning [11]. TRANSLATE learns the dependence structure among genes in the reference panel; this information is stored in the parameter estimates that are transferred to LATE for imputation of the scRNA-seq data of interest. Autoencoders have demonstrated powerful performance in other applications, such as reconstructing 2D images and 3D shapes [12]. We show with synthetic and real data that they are also powerful at imputation in highly sparse scRNA-seq data.

## RESULTS

### The LATE (Learning with AuToEncoder) Method

An autoencoder is a neural network of one or more hidden layers that allows for reconstructing the input, which is the highly sparse scRNA-seq data here, through dimension reduction, and thus generates the output with the missing values imputed (**Fig. 1A**). Each hidden layer consists of many artificial neurons (or nodes), each of which provides a certain representation of the input. An autoencoder typically contains a bottleneck layer of a lower (often much lower) dimension than that of the input, and thus achieves dimension reduction. From the input to the bottleneck layer, the salient features in the data are encoded in reduced dimensions; this half of the autoencoder is called the “encoder”. From the bottleneck layer to the output, the compressed information is gradually restored to eventually reconstruct all the values in the input; this half is therefore the “decoder”. When certain values are missing in the input, the autoencoder is therefore able to learn the dependence structure among available values and use the representations stored in the hidden layers to recover missing values.

**Figure 1:**
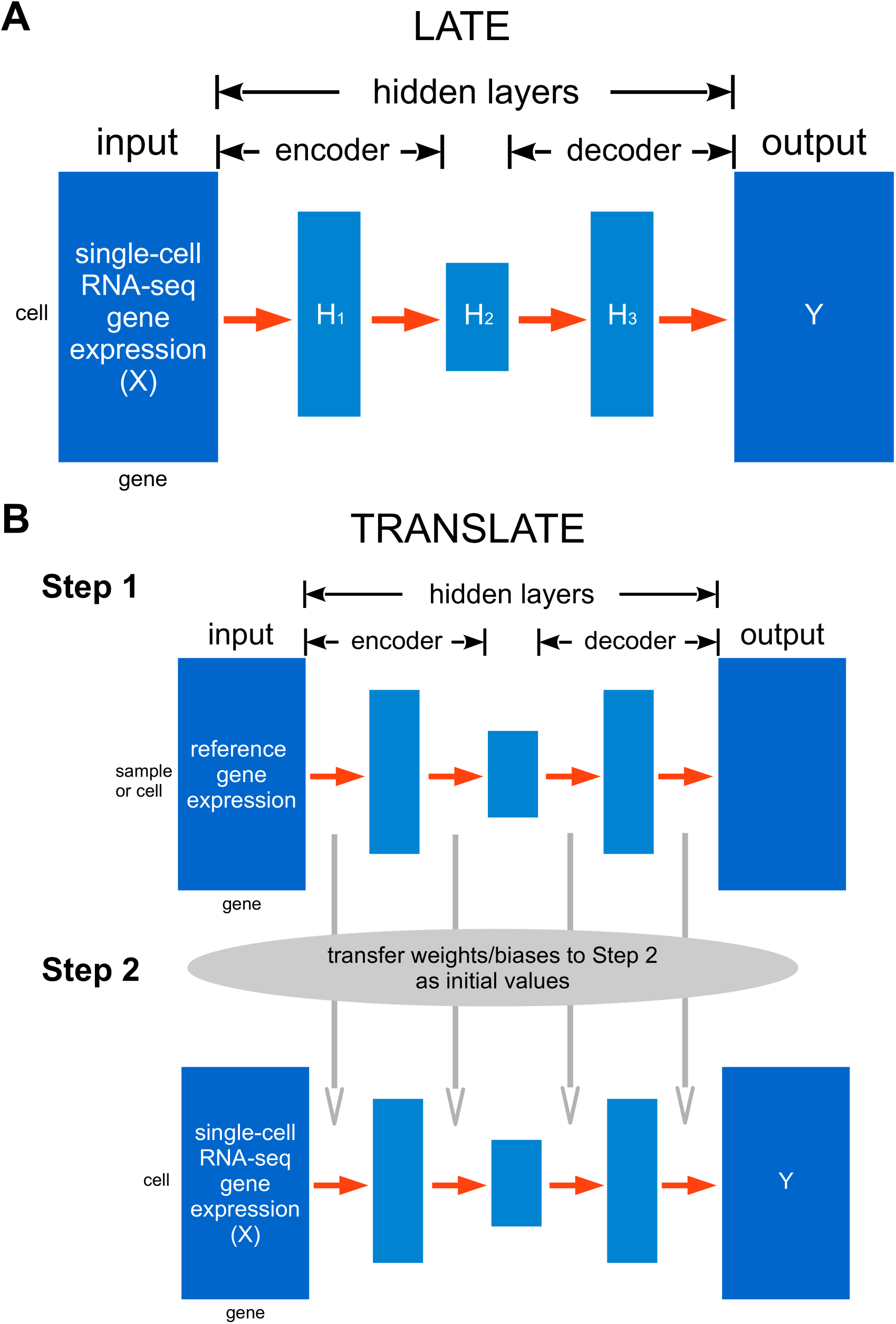
Architectures of our deep learning methods LATE and TRANSLATE for imputing zeros in scRNA-seq data. The input data matrix is represented by ***X***, and the output by ***Y***. Both matrices have the same dimensions and layout. **(A)** LATE trains an autoencoder of multiple layers directly on scRNA-seq data with random initial values of the weights and biases. **(B)** TRANSLATE incorporates transfer learning and involves two steps: the first step trains an autoencoder on a reference gene expression dataset. The weights and biases from the first step are then used to initialize the training process in the second step with the scRNA-seq data of interest as the input.

Let ***X*** be the input scRNA-seq matrix with values being log_10_-transformed read counts with a pseudocount of 1 added, i.e., log_10_ (count+1). The log_10_ transformation reduces variance in the raw read counts, which may vary from 0 to a few thousands. Let ***Y*** be the output matrix, and ***H***_*k*_ be the *k* th hidden layer. The input matrix ***X*** and output matrix ***Y*** have the same dimensions and layout. For now, we consider genes as features and cells as independent samples. Both ***X*** and ***Y*** have *m* genes (columns) and *n* cells (rows). The *i* th row in either matrix thus corresponds to the *i* th sample (cell). The first hidden layer ***H***_1_ with *l*_1_ nodes following the input ***X***_*i*_ (row vector) is derived from the following model:

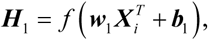

where the superscript T represents transpose, the weight matrix for the first hidden layer ***w***_1_ is *l*_1_ × *m*, and the bias term ***b***_1_ is a vector of length *l*_1_. Similarly, from the *k* th hidden layer to the *k* +1st, the model is:

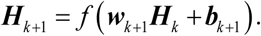

We use the rectified linear unit (ReLU) function as the activation function *f*, which means that for an arbitrary value g,

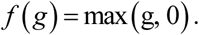

From the last hidden layer to the output layer, we have

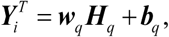

where *q* represents the last hidden layer. Our autoencoder will minimize the loss function, defined as the mean squared error (MSE) between the input and output matrix on the nonzero values:

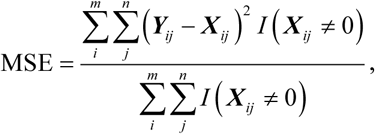

where *I* (***X***_*ij*_ ≠ 0) is the indicator function that takes on value 1 for a nonzero value in the input, and 0 otherwise.

On simulated and real data, we have experimented with 1, 3 and 5 hidden layers with 400 nodes for the single hidden layer, with 400, 200 and 400 nodes for the three hidden layers, respectively, and with 400, 300, 200, 300 and 400 nodes for the five hidden layers, respectively.

We use a suite of techniques to train the autoencoder and obtain optimal estimates of the weights and biases. Specifically, we use the backprop algorithm [13] to calculate the gradients in each layer, and the Adaptive Moment estimation (Adam) algorithm [14] to perform stochastic optimization using those gradients. We divide samples (cells) in the input data by the ratio 70:15:15 for training, validation and testing (although testing is typically not performed for imputation on the same input matrix). Optimization is performed iteratively with a randomly-generated set of parameter values as the starting point. The number of epochs (i.e., iterations) required for optimization is determined by comparing two learning curves: the curve of the MSE on the training data and that of the MSE on the validation data over epochs. We expect both learning curves to reach a minimal point with the smallest gap. If the learning curve on the training data is much lower than that on the validation data, it suggests overfitting; if both training and validation MSEs are large, it suggests underfitting. In addition, we adopt several strategies during optimization to improve efficiency and further reduce overfitting. For example, we randomly “remove” a certain percentage of the nodes in the input layer or a hidden layer, e.g., 20% or 50%. This technique, known as “dropout” [15] in deep learning, provides a computationally inexpensive but powerful method to reduce overfitting (this “dropout” technique is not to be confused with the “dropout” phenomenon in single-cell sequencing data that results in a high percentage of zeros). In each epoch, we randomly divide the training set into mini-batches (typically of 256 samples per batch), and one optimization epoch trains on multiple mini-batches until all the samples have been used for training [16].

The input data matrix for LATE treats genes as features and cells as samples, assuming that the cells are independent of one another. However, cells can be dependent as well: cells sampled from the same cell type or differentiation stage tend to be more informative of one another than cells from another cell type or another stage. Therefore, to impute the expression value of a gene in a cell, we need to account for dependence among cells in addition to that among genes. Indeed, the MAGIC (Markov Affinity-based Graph Imputation of Cells) method [5], one of the existing imputation methods for scRNA-seq data, exploits the dependence structure among cells and establish neighborhoods of cells based on the gene expression profile. Motivated by this idea, we develop the “LATE combined” method that accounts for information from both dimensions in the input data. Specifically, we perform imputation with LATE twice, first using genes as features and cells as samples, and next using cells as features and genes as samples. We obtain two imputed matrices, denoted ***Y*** ^*g*^ and ***Y*** ^*c*^, respectively, both of which have genes in the rows and cells in the columns. Next, we merge them by genes with a smaller MSE after imputation: for gene *i*, we consider the *i*th row in the two matrices, denoted by 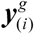 and 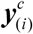, with the corresponding MSE 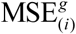 and 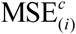, respectively. If 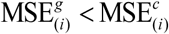, then we choose 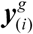. Otherwise we choose 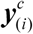. The resulting imputed matrix contains values from both matrices and has an MSE that is lower than the MSE of either ***Y*** ^*g*^ or ***Y*** ^*c*^.

### TRANSLATE: TRANSfer learning with LATE

When prior knowledge about the gene-gene relationships exists, we can incorporate this knowledge through transfer learning, which effectively runs LATE twice (**Fig. 1B**). In the first step, we train LATE on a reference gene expression data set using genes as features. In the second step, we train LATE on the scRNA-seq input data. Instead of using random values as the initial parameter values, we transfer the parameters (weights and biases) learned from the first step to the second step as initial values. As we demonstrate later, gene expression from bulk samples or from a larger single-cell sample of similar cell types may be used as the reference.

Similar to the LATE combined approach described above, we also allow for the TRANSLATE combined option, which applies TRANSLATE to the input data matrix with genes being features and applies LATE to the input matrix with cells being features. We use the same approach as in the LATE combined method to combine the two imputation matrices.

### Relationship to other scRNA-seq imputation methods

Several methods have been developed recently to impute missing values in scRNA-seq data. Some methods, such as MAGIC [5] and scImpute [3], use similarity among cells for imputation, whereas other methods, including SAVER (Single-cell Analysis Via Expression Recovery) [4], DCA (Deep Count Autoencoder) [6] and scVI (single-cell Variational Inference) [7], rely on similarity among genes. The latter methods effectively treat cells as independent samples and model the read counts in scRNA-seq data with a negative binomial distribution. Additionally, DCA and scVI take a deep learning approach and develop deep neural networks also based on autoencoders. However, whereas DCA and scVI assume read counts to follow a negative binomial distribution and estimate parameters of this distribution as part of the inference with their autoencoders, we do not make explicit assumptions on read counts. ALRA (Adaptively-thresholded Low-Rank Approximation) [17] performs randomized Singular Value Decomposition (SVD) on the gene expression matrix; whether the genes or cells are features is irrelevant with this approach. This aspect is similar to our methods that can take either genes or cells as features. Additionally, scVI accounts for batch effects in their statistical model and removes batch effects in imputation. Other methods, including ours, do not address batch effects.

### Overall imputation accuracy

We first assessed the performance between LATE and TRANSLATE, before comparing them to existing methods. We ran LATE and TRANSLATE for 1000 epochs with different architectures (with 1, 3, and 5 hidden layers) on simulated and real data (see “Generating synthetic data for assessing performance” and “Data sets without the ground truth” in Materials and Methods; the data sets are summarized in **Table S1**). We calculated the MSE, which compares the imputed data matrix to the input and the gtMSE, which compares the imputed data matrix to the ground truth, respectively (see “Assessing imputation accuracy” in Materials and Methods for the calculation of gtMSE; **Tables S2** and **S3**). Both metrics assess the deviation across genes and across cells. We observed only small differences in MSE across different architectures, and these differences typically disappeared after we fine-tuned the hyperparameters (e.g., the number of nodes in a hidden layer, the learning rate, the retain probability in the “dropout” (for deep learning) procedure, the mini-batch size, etc.). We also did not observe substantial differences between LATE and TRANSLATE in gtMSE, except on the MAGIC mouse data. Compared to LATE, TRANSLATE reduces both MSE and gtMSE by half on the MAGIC mouse data, indicating that the reference data significantly improves imputation. However, the reference and the input in the MAGIC mouse data come from the same data set, which means that the dependence structure is essentially identical in the reference and in the input. In reality, however, such perfect reference is usually not available.

We next compared our methods with other imputation methods, namely DCA, scVI, SAVER, MAGIC, ALRA and scImpute, and summarized MSEs and gtMSEs in **Table S3**. We used the default settings of these methods wherever possible. scImpute requires the number of clusters (cell types) as part of the input. We used the true number of clusters when known and the value 1 otherwise. Since TRANSLATE uses the reference data, which can contain a substantial amount of information unavailable to other methods, it is unfair to compare other methods to TRANSLATE. Instead, we compare other methods only with LATE here.

Our LATE methods achieve a lower MSE (with respect to the input) than other methods in most cases. When the ground truth is available, our methods have a lower gtMSE than the other methods on all the synthetic data sets except for the GTEx_4tissues data. Since MSE and gtMSE are calculated across genes and across cells, if imputed values for a gene were allocated to the wrong cells, the MSE (and gtMSE) would be larger. We can confirm this by calculating the Pearson correlation across all the genes between the imputed values and the ground truth for each cell and generating the histogram of all the cells. The histograms from our methods are generally closer to 1 than that of other methods (see an example in **Supplementary Fig. S1** between LATE and MAGIC).

The GTEx_4tissues data set was generated from the data in the Genotype-Tissue Expression (GTEx) consortium. Note that we calculated gtMSE_all_ (using all values) for GTEx_4tissues, but gtMSE_nz_ (using only nonzero values) for other data sets. For GTEx_4tissues, DCA achieves the lowest gtMSE, followed by our methods, scImpute, MAGIC, scVI, ALRA and SAVER. Recall that both DCA and scVI estimate the “dropout probability” of an expression value of zero is due to scRNA-seq dropout and impute the zeros only when they have a high dropout probability. The MSE/gtMSE comparison above indicates that DCA can be successful at detecting true zeros and is more successful than scVI. On the other hand, the DCA model for nonzero values may be biased. Note that for this data set, SAVER achieved the lowest MSE, but the highest gtMSE (20 times higher than LATE did), suggesting that the SAVER model is overfitting the nonzero values in the input and does not extrapolate well for zeros. Also note that for all the data sets, scImpute achieved an MSE of zero or nearly zero but higher gtMSE than our methods. scImpute performs imputation cell by cell, and divides genes of each cell into genes with credible expression values and those needing imputation. The higher gtMSEs from scImpute suggest that this approach does not fully capture the complex dependence among genes and cells.

We can further break down gtMSE_all_ by calculating of contributions from different sources: gtMSE_nz_, the gtMSE for the nonzero values in the input; gtMSE_biol_, the gtMSE for the zeros in the ground truth, which represent no expression and therefore are termed “biological zeros”; and gtMSE_tech_, the gtMSE for the zeros introduced into the input by masking nonzero values in the ground truth – these values are termed “technical zeros” (see “Assessing imputation accuracy” in Materials and Methods; **Supplementary Fig. S2**; **Table S4**). Specifically, the ground truth for the MAGIC_mouse data does not contain biological zeros. Only gtMSE_tech_ can be calculated and its value does not differ much from the corresponding gtMSE_all_ of each method under comparison. The ground truth for the GTEx_4tissues contains 50.20% of biological zeros. LATE, TRANSLATE, DCA and scVI all have a higher gtMSE_biol_ than gtMSE_tech_, whereas SAVER, MAGIC, ALRA and scImpute are the opposite. On this data set, DCA achieved lower gtMSE_all_ than our methods, which may be explained by its much lower gtMSE_biol_ and slightly higher gtMSE_tech_ than ours. The two PBMC data sets (PBMC_G949 of 949 genes and 21K cells, and PBMC_G949_10K of the same 949 genes but from 10K cells) were generated from subsets of a large, unimputed scRNA-seq data set with additional masking. The ground truth for these two data sets contains 51.10% and 67% of zeros, respectively. These zeros are a mixture of biological and technical zeros that cannot be distinguished, and gtMSE_biol_ thus refers to the error on this mixture. For these data sets, gtMSE_nz_ is the metric for the overall performance, and gtMSE_tech_ reflects the error on the additional technical zeros that we introduced to the input through masking. Our methods and ALRA infer a much lower gtMSE_tech_ than gtMSE_biol_, whereas other methods infer a gtMSE_tech_ that is similar to slightly higher than gtMSE_biol_. Furthermore, gtMSE_tech_ of our methods is substantially lower than that of any other methods, which explains the low gtMSE_nz_ from our methods. In summary, our methods impute technical zeros much better than other methods, whereas DCA, which has a competitive performance, infers biological zeros better than our methods.

On the four real scRNA-seq data sets with no ground truth (see “Data sets without the ground truth” in Materials and Methods), we calculated only MSE to compare the performance of different methods (**Table S3**). These data sets are derived from the large scRNA-seq data generated by 10x Genomes in human peripheral blood mononuclear cells (PBMCs) and in mouse brain. LATE and TRANSLATE produced similar MSEs, both being no more than a third of the MSE from scVI, SAVER and MAGIC and just 2% of the MSE from ALRA. DCA achieved 41% of the MSE by LATE combined on the PBMC_G5561 data, but 450% of the MSE by LATE combined on the PBMC_G9987 data in which the number of genes nearly doubles. scImpute was unable to run on these two PBMC data sets. Due to similar performance between LATE and TRANSLATE, we only ran LATE on the two mouse brain data sets. Other methods either took a much longer time or were unable to run on these two data sets (see detail in “Scalability” in Results).

### Recovery of nonlinear gene-gene relationships

We expect a good imputation method to perform well not only on the overall accuracy, but also in capturing the true dependence structure among genes. The previous section shows that a small gtMSE indicates that the imputation is close to the truth in all aspects. A small MSE, which is calculated only on the nonzero values in the input, does not necessarily represents good performance. It is of interest to investigate whether the gtMSEs from our and other methods are small enough to recover gene-gene relationships. Additionally, when assessing pairwise relationships, linear relationships between genes are generally easy to capture, whereas nonlinear relationships are much more difficult. Here, we examine the performance of our methods and other methods on recapitulation nonlinear relationships in gene pairs.

The synthetic data sets we have generated contain different types of nonlinear relationships. In the MAGIC_mouse data, the diffusion process in the MAGIC method provides an approach to generate gene expression profiles that evolve over diffusion time and that enhance the correlation structure over diffusion time [5]. As the number of diffusion time points, denoted by *t*, increases, the diffusion process produces more sharply defined nonlinear patterns. Although the imputation results from MAGIC are less satisfactory, their model using a diffusion process allows us to generate synthetic single-cell data with sharply defined nonlinear patterns (see such pattern in the ground truth in **Fig. 2**; by contrast, gene-gene relationships from other synthetic data sets are more similar to having multiple clusters; see below). The step of masking down to a nonzero rate of 10% removes most of the nonlinear features in the input, although the upper “arm” remains (**Fig. 2** input). Both LATE and TRANSLATE recovered the nonlinear relationship. Surprisingly, even though the ground truth and input were generated by MAGIC, it could not recover this pattern. None of the other methods recovered this pattern, either.

**Figure 2:**
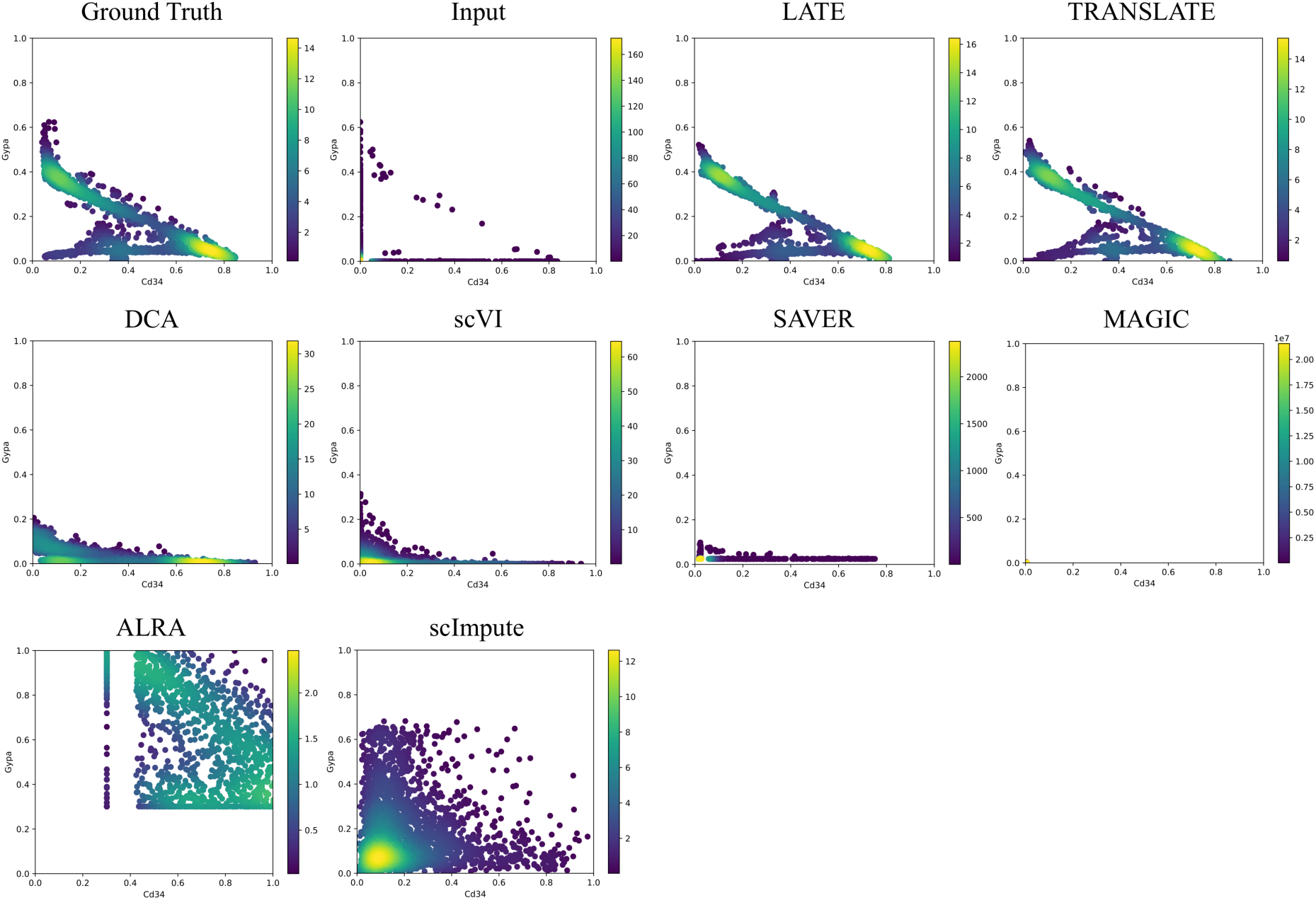
Two genes (*Cd34* vs *Gypa*) with a nonlinear relationship in the synthetic data based on the mouse bone marrow data (MAGIC_mouse; 16,114 genes and 2,576 cells). Each dot is a single cell. The color bar indicates the Gaussian kernel density estimates of data points. The input contains only 10% of the values in the ground truth. Methods under comparison are: LATE, TRANSLATE, DCA, scVI, SAVER, MAGIC, ALRA and scImpute. Only LATE and TRANSLATE recovered the nonlinear relationship. Data points for ALRA are mostly outside the plotting area. The full view is in **Supplementary Fig. S3**. The data points in the results from MAGIC are concentrated near the origin (**Supplementary Fig. S4**). See three additional gene pairs with different types of nonlinear relationship in **Supplementary Figs. S5-S7.**

The other type of nonlinear patterns may be better described as forming multiple clusters (the ground truth in **Figs. 3** and **4**). Recall that the ground truth for the GTEx_4tissues data set (**Fig. 3**) is the partial GTEx data, which are bulk gene expression values, and that the ground truth for the PBMC_G949 data set is the scRNA-seq data in the PBMCs with a nonzero rate of 41% (**Fig. 4**). The nonlinear patterns in these data sets are therefore directly obtained from the real data. Since the data were not collected over time, it is reasonable that these nonlinear patterns may reflect subpopulations of cells, rather than a trajectory over which the cells evolve.

**Figure 3:**
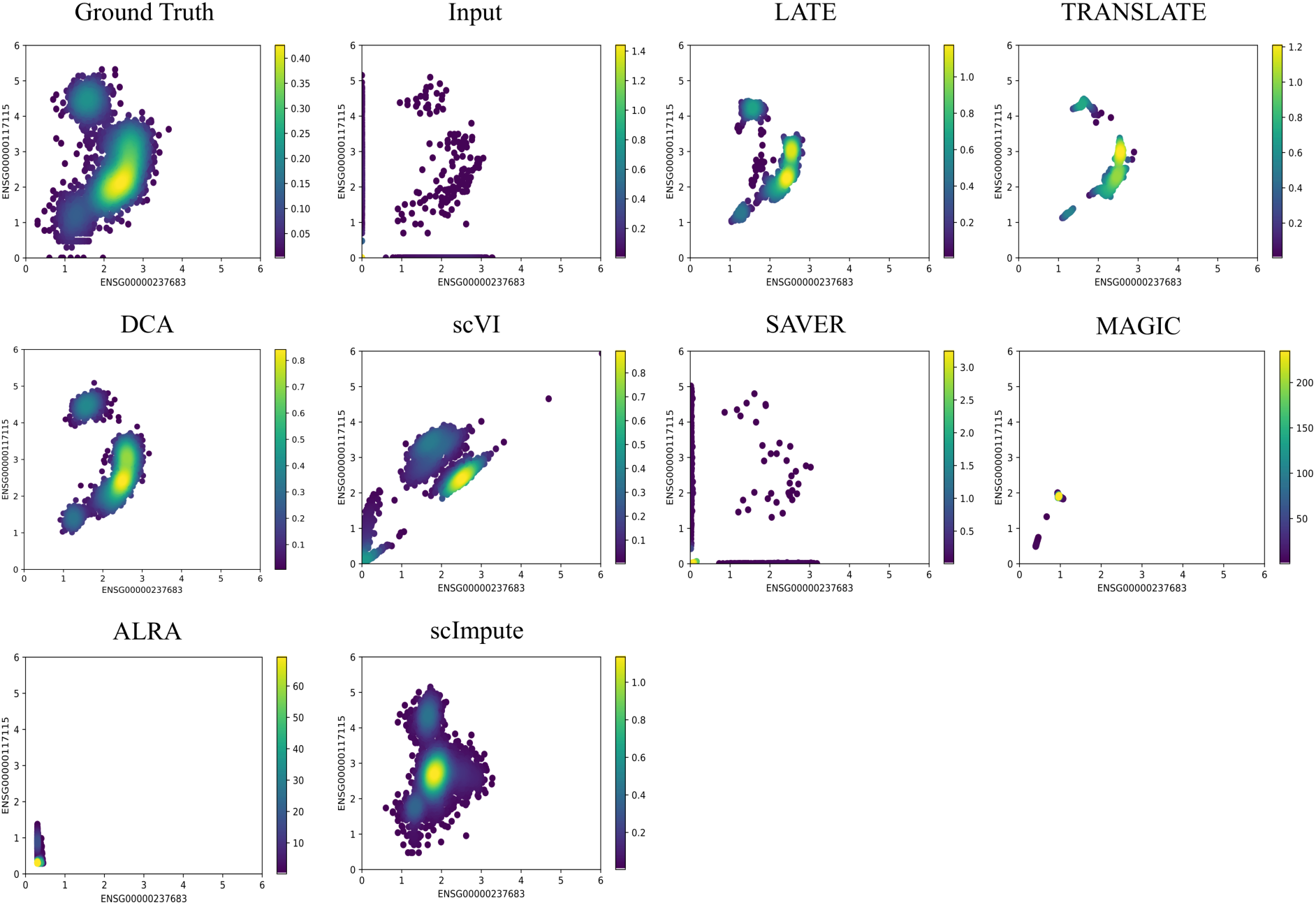
Two genes (*ENSG00000237683* vs *ENSG00000117115*) with a nonlinear relationship in the synthetic data based on the GTEx (Genotype-Tissue Expression) consortium (GTEx_4tissues; 56,202 genes and 3,164 samples). Each dot is a single cell. The color bar indicates the Gaussian kernel density estimates of data points. Methods under comparison are: LATE, TRANSLATE, DCA, scVI, SAVER, MAGIC, ALRA and scImpute. LATE, TRANSLATE, DCA and scImpute recovered the nonlinear relationship reasonably well. See additional gene pairs with different types of nonlinear relationship in **Supplementary Figs. S8-S10**.

**Figure 4:**
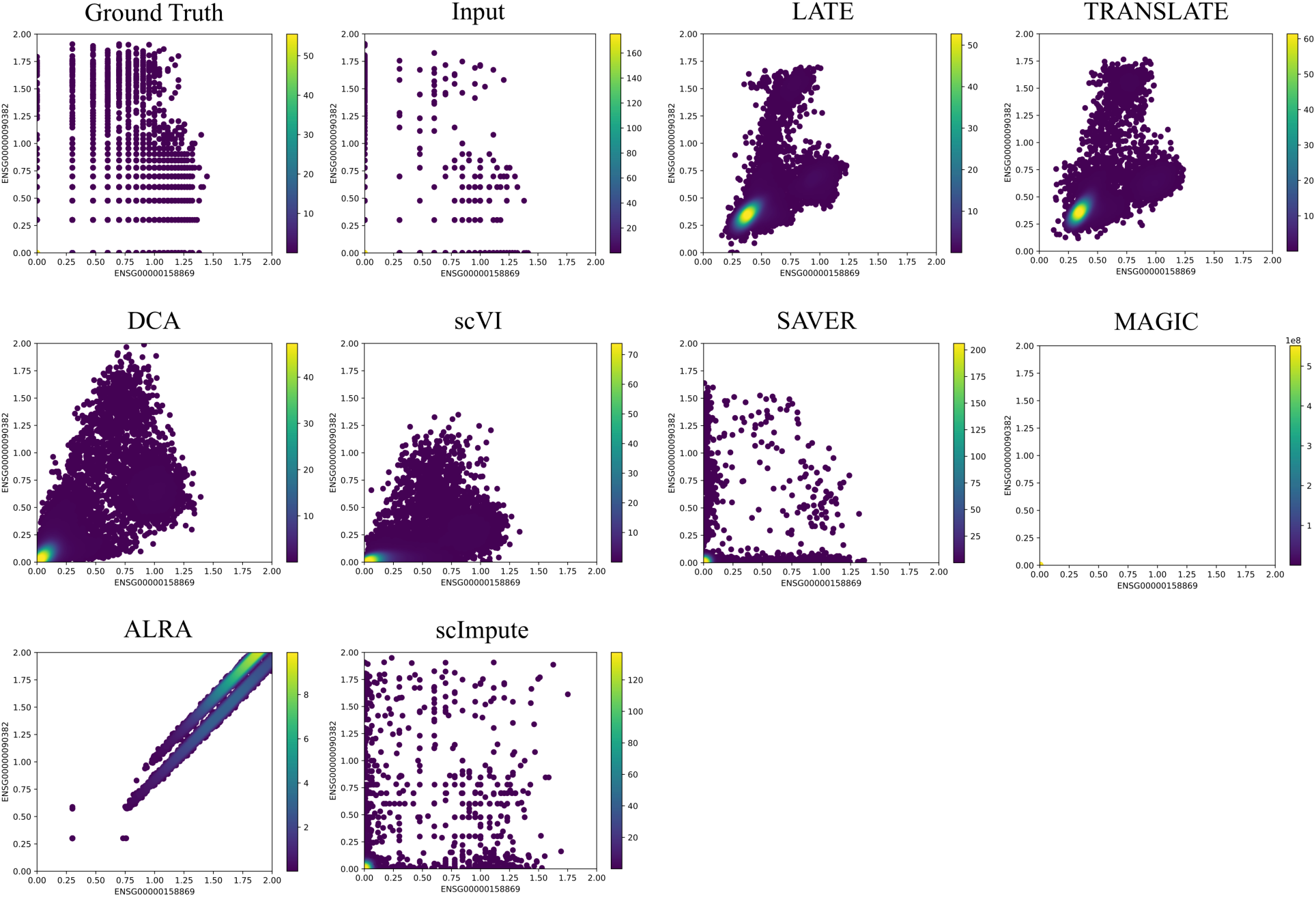
Two genes (*ENSG00000158869* vs *ENSG00000090382*) with a nonlinear relationship in the small synthetic data based on the 10x Genomics PBMC data (PBMC_G949; 949 genes and 21,065 cells). Each dot is a single cell. The color bar indicates the Gaussian kernel density estimates of data points. Methods under comparison are: LATE, TRANSLATE, DCA, scVI, SAVER, MAGIC, ALRA and scImpute. LATE, TRANSLATE, DCA and scVI all recovered the nonlinear relationship reasonably well, although scVI inferred a much-reduced range. Similar to **Fig. 2**, the data points from MAGIC are also concentrated near the origin (**Supplementary Fig. S4**). See additional gene pairs with different types of nonlinear relationship in **Supplementary Figs. S11-S12.**

On the input data derived from these ground truth data sets, LATE and TRANSLATE continue to recover the truth on selected gene pairs (**Figs. 3** and **4**). The performance of DCA and scVI improves drastically, although not necessarily better than our methods. These improved results suggest that DCA and scVI may be sensitive to how the data are generated. Unlike DCA and scVI that model the read count of genes with negative binomial distributions, MAGIC focuses on the dependence structure among cells and does not model read counts. If the single-cell data were generated by the MAGIC, which is likely incompatible with the model behind DCA or scVI, the latter two methods may not be able to produce sensible results. The performance of MAGIC also improves somewhat, although it still produced patterns that capture only partial features of the true relationships. SAVER produces scatterplots that are similar to the input, confirming our earlier observation that its imputation runs the risk of overfitting to the nonzero values in the input. ALRA is unable to recover either pattern. scImpute performs well on GTEx_4tissues but not on PBMC_G949.

Additional gene-gene relationship plots on other data sets without the ground truth are in **Supplementary Figs. S13-16**. Although only a few gene pairs are shown in the main text and supplementary figures, the performance of the methods is largely consistent with their MSE and performance on other data sets.

### Separation of cell types

Single-cell data often consist a mixture of cell types, and the high dropout rate tends to obscure the differences among cell types. While broad classifications of cell types are still possible with the aid of marker genes, the obscured relationship makes it difficult to identify novel or poorly characterized cell types. Imputation is therefore a powerful approach to help recover cell types: it provides higher-quality data for downstream clustering analyses, which separate known cell types and may suggest novel ones. Using single-cell data with known cell type labels, we can examine imputation performance on separating cell types.

We applied all the methods discussed above to the PBMC_G949_10K data, which was derived from a larger PBMC data set with cell type labels, generated tSNE (t-Distributed Stochastic Neighbor Embedding) plots of all the cells before and after imputation, and colored the cell types (see “Principal Component Analysis (PCA)-based tSNE plots for visualization” in Materials and Methods; **Fig. 5A**). We also calculated the between-cluster sum of squares (BCSS) to quantify the separation of the cell types and the within-cluster sum of squares (WCSS) to measure the closeness of cells to each other of the same cell type (see “Metrics for cell type separation” in Materials and Methods; **Fig. 5B**). When running TRANSLATE, we used the other 30K cells from the 40K cells with cell type labels as the reference, which we refer to as reference i) (**Table S2**). Indeed, whereas cells of the same type (or color) are largely grouped in a cluster in the tSNE plot of the ground truth, different cell types are completely scrambled and indistinguishable in the tSNE plot of the input, resulting in both WCSS and BCSS being low. LATE, TRANSLATE, DCA and scVI all managed to group similar cells and separate cell types after imputation, producing a tSNE plot similar to that of the ground truth and achieving a high BCSS and a low WCSS. By contrast, none of the other methods separates the cell types: SAVER produced a tSNE plot that is almost identical to the input with similarly low BCSS and low WCSS. scImpute did not separate the cell types, either, but achieved a higher BCSS than SAVER. MAGIC and ALRA separate cells into tiny clusters that do not reflect the cell types, also indicated by their low BCSS and high WCSS.

**Figure 5.**
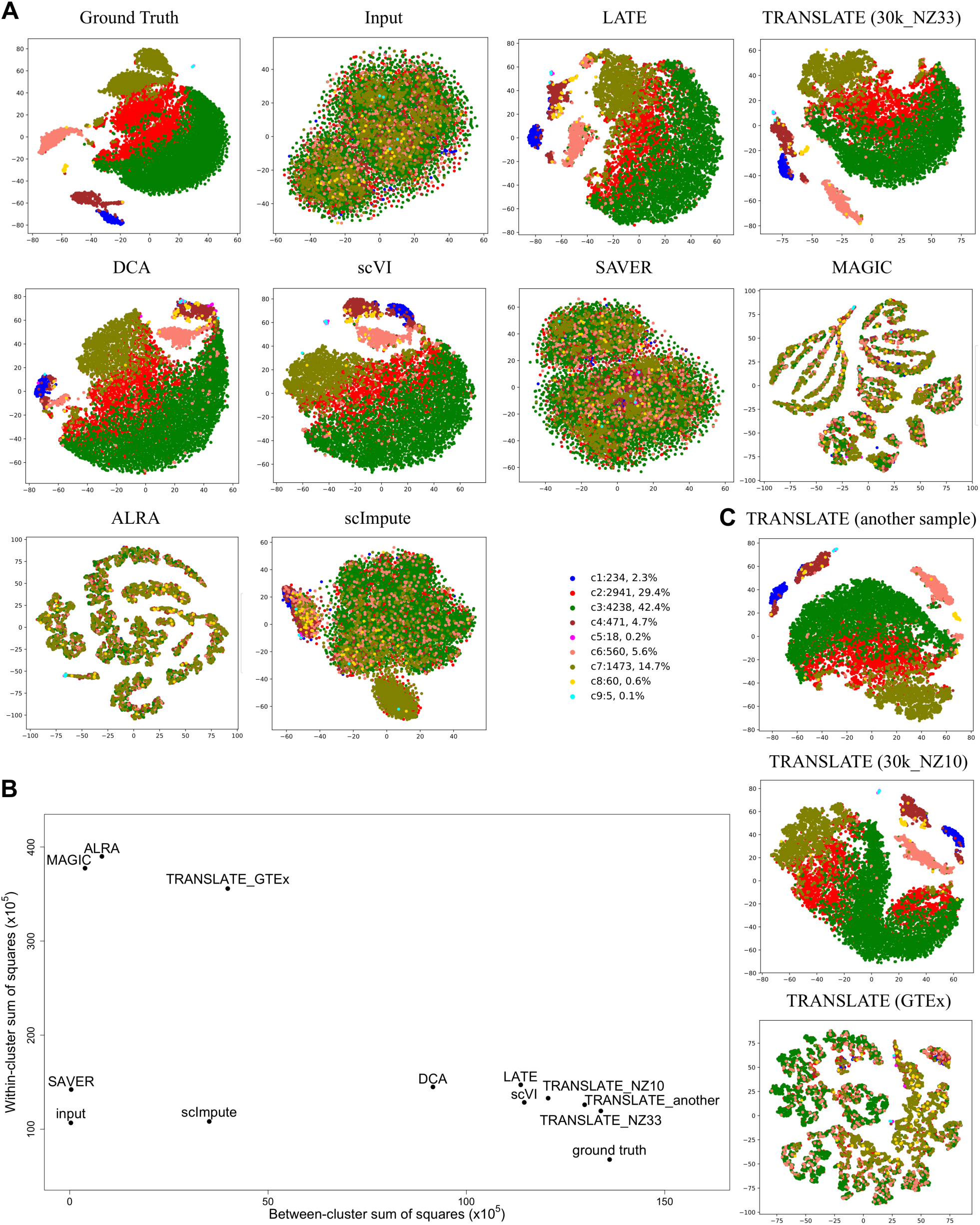
tSNE plots of cells from the synthetic data based on the 10x Genomics PBMC data with known cell types (PBMC_G949_10K; 949 genes and 10K cells). Each dot is a single cell, and each color indicates a cell type. Methods under comparison are: LATE, TRANSLATE, DCA, scVI, SAVER, MAGIC, ALRA and scImpute. **(A)** Comparison of all the methods. LATE, TRANSLATE (30k_NZ33 as the reference), DCA and scVI all recovered the clustering. **(B)** Between-cluster and within-cluster sum of squares to assess the cell type separation. **(C)** Comparison of different reference gene expression data sets in TRANSLATE. Using other scRNA-seq data as the reference (e.g., another sample, and 30k_NZ10), TRANSLATE also managed to separate rare cell types (c8 in yellow, c5 in magenta and c9 in turquoise). The bulk gene expression of multiple tissues from GTEx did not serve as a good reference for separating cell types.

A closer look at the results from LATE, TRANSLATE, DCA and scVI reveals subtle differences: apart from the larger clusters being visually separated, TRANSLATE with the 30K PBMC single-cell data being the reference (the 30k_NZ33 data set, which has a nonzero rate of 33%) manages to also separate the c8 cluster of 60 cells (0.6% of the total 10K cells; represented by yellow dots in **Fig. 5A, C**) and the even smaller clusters of c5 (18 cells; magenta in **Fig. 5A, C**) and c9 (5 cells; turquoise in **Fig. 5A, C**) from the larger clusters, although c5 and c9 are not distinguishable. Since tSNE generates a different layout each time it is run, we ran tSNE again on the TRANSLATE imputed data multiple times, and c8 and c9 are always separated from larger clusters. In addition to reference (i), we next experimented with three other references for TRANSLATE in order to understand the role of reference in the cell type separation (**Fig. 5C**; **Table S2**): (ii) “another sample”: the single-cell PBMC data from another sample collection of the same individual; (iii) “30k_NZ10”: the 30K PBMC cells with masking (the nonzero rate is also 10%); and (iv) all the tissues in the GTEx data. Most cells in c8 and all the c5 and c9 cells are clearly clustered together and separated from other clusters in the tSNE plots from references (ii) and (iii), whereas using GTEx as the reference does not lead to satisfactory results (**Fig. 5B, C**). However, the MSEs do not differ too much among the four references: both MSE and gtMSE are only slightly higher under the GTEx reference than the other three references (**Table S3**). The poor performance with the GTEx reference on separating cell types is likely due to the tissue types in the GTEx data being very different from the PBMCs. As a result, the weights and biases TRANSLATE learns from GTEx are a rather poor starting point for the single-cell input data. Among other methods we compared with, scVI and LATE can separate most of c5 and c9 cells, but cannot group the c8 cells together. DCA is not able to group the c8 or c9 cells together or separate them from other clusters. This analysis indicates the additional advantage of having a relevant reference in our TRANSLATE method, and also provides an example where a reference from irrelevant tissue or cell types may increase the MSE and produce less satisfactory imputation results.

The synthetic data above test the performance of the imputation methods in realistic settings. For comparison, we also simulated scRNA-seq read count data of 1,000 genes, 20,000 cells from six clusters using the R package Splatter [18] (see “Simulation with the R package Splatter” in Materials and Methods). The statistical model underlying this simulation procedure assumes the same relative magnitude of a gene (to other genes in the same cell) across cells. The dependence among genes in a cell is therefore similar to that in a multinomial distribution: for a large number of genes, they are nearly independent of each other within a cell. However, these assumptions are far from reality, as genes are known to act in complex regulatory networks. None of the methods under comparison is able to recover the clusters (**Supplementary Fig. S17**) due to the weak dependence among genes. However, their performance on this data set is not indicative of that on real and more complex data. This is the reason why we generated other synthetic data from real gene expression data, such that the complex regulatory relationships among genes were retained.

We next examined the imputation performance of our methods on data with batches. We reanalyzed the mouse retina data previously analyzed by scVI [19] (see Data Availability; **Fig 6**). This data contains 13,166 genes and 19,829 Cells with a nonzero rate of 6.7%. The tSNE plot on the input of log10-transformed count data displays the claw-shaped clusters of cell types; the claws reflect the two batches in this data set. scVI provides a cleaner separation than our LATE method does, although the claw shapes remain in several cell types even though scVI has removed the batch effect. This is particularly encouraging to us as our methods do not account for batch effect in the deep learning architecture. The tSNE plot indicates that LATE still manages to group similar cell types together while retaining the batch effect. For downstream analysis, one may apply batch correction (e.g., [20]) to LATE-imputed data.

**Figure 6.**
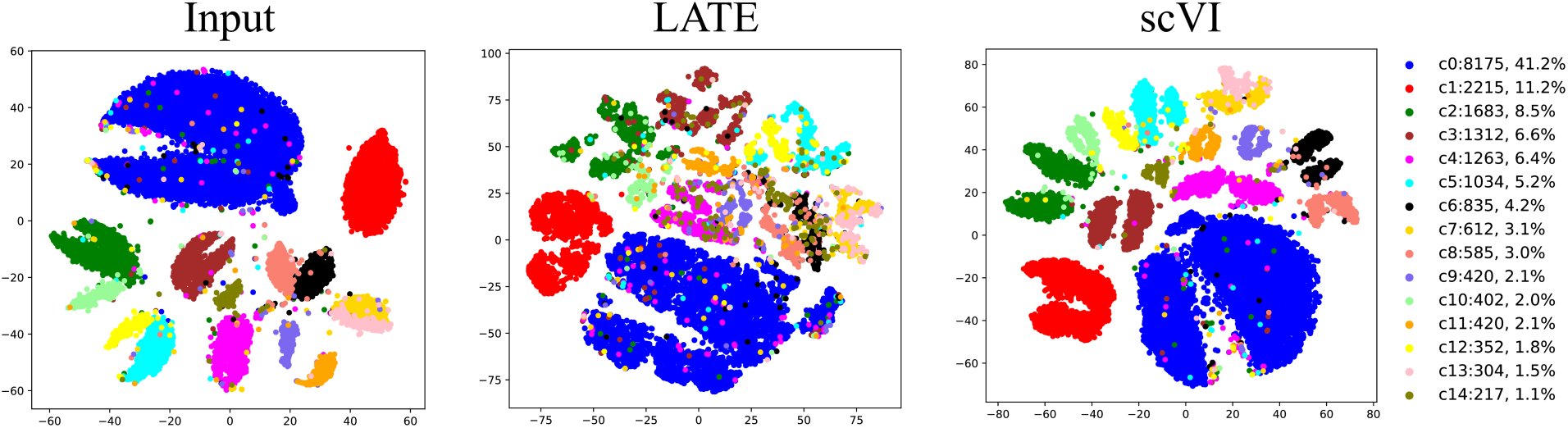
Imputation performance on cell type separation in the mouse retina data with batch effects. A tSNE plot is generated for the input, for the output from LATE, and for the output from scVI. The claw-shaped clusters are due to two batches in the data.

### Scalability

We used the mouse brain data also from 10x Genomics to test scalability (see “Data sets without the ground truth” in Materials and Methods). The entire mouse brain data consists of 28K genes and 1.3M cells, with a nonzero rate of 7% (mouse_brain_28K). We ran LATE but not TRANSLATE on this large data set because LATE and TRANSLATE have similar performance on most data sets, and because our interest here is primarily scalability. Nevertheless, TRANSLATE in principle can run on this data set as well, as it is essentially running LATE twice. Note that although LATE and TRANSLATE provide the “combined” option, this option does not work on this data set. This is because this option first uses genes as features and next uses cells as features. However, our code based on TensorFlow currently cannot handle 1.3M features, which is the number of cells, on existing Graphics Processing Units (GPUs) or Central Processing Units (CPUs) even with a large Random Access Memory (RAM).

To further investigate the scalability of LATE, we generated multiple data sets from the mouse brain data, by sampling the cells from the data set of 10K genes and that of 28K genes. We ran LATE for 300 epochs on these data sets on multiple GPUs (including NVIDIA GTX 1080 Ti, V100, and Titan RTX GPUs) and recorded the runtime (**Fig. 7; Table S5**). We used 300 epochs here as this is the default setting in DCA, the most competitive method. Although several models of GPUs were used in this investigation, the runtime is essentially independent of the GPU models. The increase in the runtime increase is better than linear with respect to the number of cells or the number of genes. Specifically, LATE took only less than 11 hours to impute the entire mouse brain data (28K genes and 1.3M cells), and 7.3 hours to impute half of the genes (also 1.3M cells).

**Figure 7.**
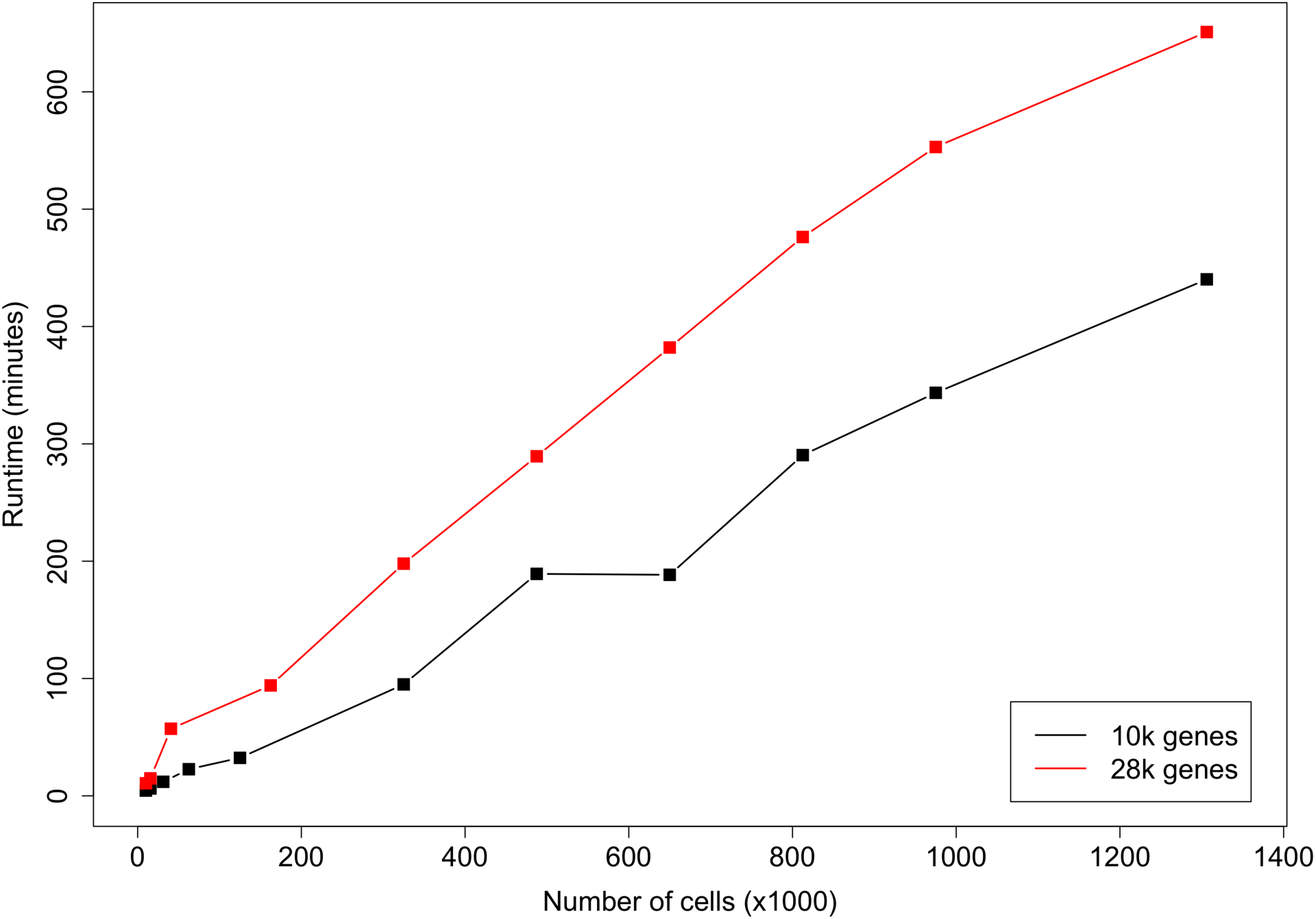
GPU runtime of LATE (genes as features) on multiple subsets of the mouse brain scRNA-seq data from 10x Genomics. The entire data set contains 28K genes and 1.3M cells. We sampled a subset of 10K genes. For the 28K and 10K genes, we further sampled multiple subsets of cells and used them as the input.

We further ran the data sets of 10K genes on a CPU (Dell M820 Blade with 1TB RAM); see **Supplementary Fig. S18 and Table S6**. Similar to the trend for GPUs, the increase in runtime is also linear or better with respect to the number of cells. Additionally, LATE took 53.2 hours on this CPU to impute the 1.3M cells, which is over seven times (53.2/7.3=7.3) the corresponding GPU time. In general, the CPU runtime is at least seven times the GPU runtime. We ran other imputation methods on this CPU for the data set of 10K genes and 1.3M cells. DCA took 42 hours for the default of 300 epochs, scVI completed only 8% of training after running for 56 hours, MAGIC and ALRA failed to run, and SAVER reported memory error after running for more than 129 hours. scImpute failed to run on this data set, and also took a much longer time than other methods on smaller data sets. For example, scImpute took 68 hours to run on a subsample of 15K cells from mouse_brain_10K (10K genes and 15K cells), and kept running after 6 days on a subsample containing 31K cells (10K genes and 31K cells), and failed to run on a subsample containing 62K cells (10K genes and 62K cells; half of the mouse_brain_10K data).

## DISCUSSION

Here we present our novel deep learning algorithms for efficiently and accurately imputing zeros in highly sparse single-cell RNA-seq data that often have a low nonzero rate. Our algorithms are nonparametric: they do not make assumptions on the statistical distributions of the single-cell data. Similar results between LATE and TRANSLATE on most of the data sets examined here indicate the effectiveness of autoencoders in quickly reaching optimality or near optimality, regardless of the starting point. Our algorithms are also fast and highly scalable.

Additional work is needed to improve our methods. Here we manually tuned hyperparameters, but did not explore the possibility of systematically tuning these parameters, as that implemented in DCA. When combining imputation results from both dimensions, our current approach of combining two imputed data matrices is rather ad hoc; joint optimization for both dimensions simultaneously may help further improve the results.

In our simulation study, we reported the results from generating the input data by masking 90% of the values in the ground truth, which corresponds to assigning a Bernoulli distribution of probability 0.9 to each value in the data matrix. We also experimented with subsampling from the ground truth with other distributions to also arrive at a nonzero rate of 10%, such as an exponential distribution (similar to that used in van Dijk et al. [5] for MAGIC) and a binomial distribution (similar to that used in Lopez et al. [7] for scVI). However, the performance of the methods we investigated here varied with different sampling distributions (see [21] for a recent exploration of the statistical properties of imputation with principal component analysis under heterogeneous missingness). Since it is unknown which is the true process during dropout, we reasoned that the sampling distribution in simulation should be simple and introduce minimal additional uncertainty in the assessment of the performance.

In LATE and TRANSLATE we make the central assumption that all the zeros in the scRNA-seq data are missing values. As a result, our methods impute zeros always with a nonzero value. However, it is desirable to distinguish biological zeros from technical ones. One way to approach this problem is to estimate the probability of an expression value is a dropout event. Both scVI and DCA have an explicit model for the read counts and indeed estimate this probability, although their performance is rather unsatisfactory based on our comparison. DCA infers biological zeros well (low gtMSE_biol_) but cannot impute technical zeros well (high gtMSE_tech_), whereas scVI does not do well on either type of zeros. It is therefore unclear whether the unsatisfactory performance in DCA and scVI is due to scRNA-seq data not containing enough information for distinguishing the zeros, or whether these two methods being unable to make such distinction. Our method comparison on simulated and real data indicates that our nonparametric approach to imputation based on an autoencoder achieves satisfactory performance overall, which is driven by the superior performance on technical zeros. Work is still needed to better handle biological zeros without sacrificing the accuracy on technical zeros.

## MATERIALS AND METHODS

### Assessing imputation accuracy

We calculate the MSE with respect to the nonzero values in the input data matrix as an overall measure of imputation accuracy. When the ground truth ***T*** is available, which has the same dimension as ***Y***, we can also calculate another MSE with respect to all the values in ***T*** in order to assess how close the imputed data matrix is to the ground truth:

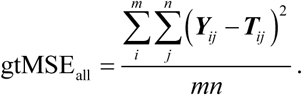

There are cases where zeros in the ground truth may still represent missing values: for example, we may use a real single-cell data set with a high nonzero rate as the ground truth and mask certain values to generate synthetic input. Then the zeros in the ground truth still represent missing values and should not be interpreted as no expression. In such cases, it is sensible to exclude the zeros when calculating this MSE:

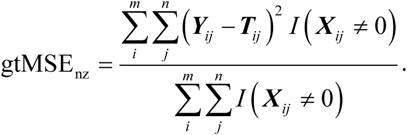

We can further break gtMSE down to that for biological zeros and that for technical zeros. Biological zeros are zeros in the ground truth and indicate no expression. Technical zeros are entries that have nonzero values in the ground truth but zeros in the input; in other words, these zeros are due to technical artifacts (such as dropouts) and do not reflect the actual expression levels (**Supplementary Fig. S2**). Let *p*_0_ be the percentage of zeros in the input, and *q*_0_ the percentage of zeros in the ground truth. Then *q*_0_ is the rate of biological zeros, whereas *p*_0_ − *q*_0_ is the rate of technical zeros. Let gtMSE_biol_ be the gtMSE for biological zeros and gtMSE_tech_ for technical zeros. Recall that there are *m* genes and *n* cells. To calculate these gtMSEs, we can break down the total sum of squares in gtMSE_all_ in two ways:

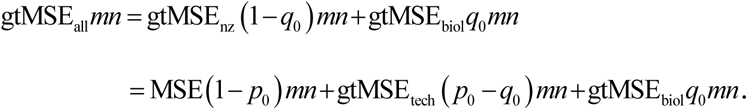

Cancelling the term *mn*, we have

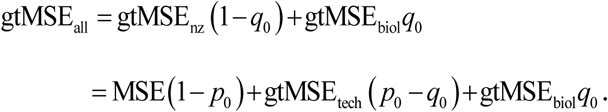

Then we have

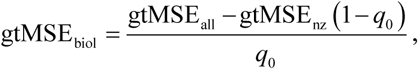

and

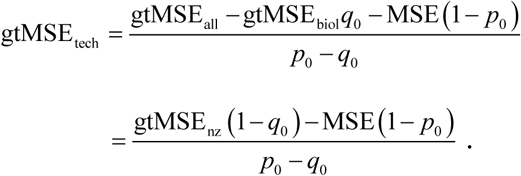

In the special case where the ground truth has no biological zeros (i.e. *q*_0_ = 0), gtMSE_all_ = gtMSE_nz_, and we can compute only gtMSE_tech_:

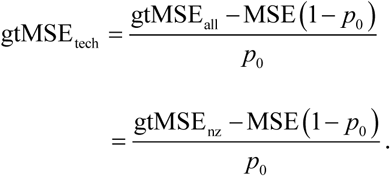

### Generating synthetic data for assessing performance

Using real gene expression data sets (from bulk tissues and single cells) in human and mouse (see Data Availability), we generated several synthetic data sets, including the input and the corresponding ground truth, for assessing the performance of imputation methods (summary statistics of the data sets in **Table S1**).

1. The MAGIC_mouse data. The authors of the MAGIC method analyzed a mouse bone marrow data set of 16,114 genes and 2,576 cells [5, 22]. This data set contains cells at multiple differentiation stages of hematopoiesis; each stage had been associated with the expression of a number of surface marker genes. This data set therefore contains dynamic changes of gene expression and is valuable for assessing how the imputation method captures the dynamics. The original scRNA-seq data is highly sparse. To enhance the dynamic changes in gene expression and the correlation structure among genes, we applied the MAGIC method to generate the imputed data set. Although the dynamic patterns produced in the imputed data set may not be real, the imputed data provide the ground truth that contains sharply-defined nonlinear patterns. We randomly selected 30% of the cells in the imputed data and split the data into two subsets. The subset containing 30% of the cells was used as the ground truth. We then generated the input data by randomly masking 90% of the values in the ground truth as zero. We applied LATE and other methods to the input and compared the imputed data matrix with the input and with ground truth. We used the other subset that contained 70% of the cells as the reference expression data for TRANSLATE.
2. A subset of the GTEx data from human (GTEx_4tissues). The GTEx consortium contains bulk gene expression data from 53 tissues (GTEx Portal v7), which is useful for us to assess the impact of using relevant and irrelevant tissues as the reference in TRANSLATE. This data set also contains many zeros that are more reliable indication of no expression, which we may use to assess how the imputation method perform in the presence of true zeros. From the 53 tissues, we selected four tissues with large sample sizes as the ground truth: muscle (564), heart (600), skin (1,203) and adipose (797). This data contains 56,202 genes from 3,164 samples, with a nonzero rate of 49.8%. We then randomly masked 90% of the values as zero to create an input data set. When assessing the performance of TRANSLATE, we considered two references: i) the entire GTEx data from all the tissues, including the four tissues in the input; and ii) the GTEx data without the four tissues in the input. We were interested in whether including the right tissues in the reference is necessary for imputation. Note that when calculating gtMSE for GTEx_4tissues, we needed to calculate gtMSE_all_, instead of gtMSE_nz_, that were used for other data sets. This is because zeros in the GTEx data were obtained from bulk sequencing and therefore should be interpreted as no expression.
3. A small subset of the data in human PBMCs from 10x Genomics [23] (PBMC_G949). The original data set contains 68K cells. We selected 949 genes in 21,065 cells, such that the total read count per gene is >10K and the total read count per cell is >1.5K. This data set has a nonzero rate of 41% and was used as the ground truth (zeros were discarded when calculating gtMSE). We randomly masked nonzero values so that the resulting data contains only 10% of nonzero values. When applying TRANSLATE, we explored several options for the reference, which is explained in detail in Results.
4. Another subset of the 10x Genomics PBMC data with cell type labels (PBMC_G949_10k). To assess the ability of the methods in separating different cell types, we generated another data set from the same PBMC data as mentioned above. The original data set contains 40K cells with cell type labels and 10 cell types. We focused on the same 949 genes as in the PBMC_G949 data set, subsampled 10K cells while retaining the relative frequencies of the ten cell types; this data set, with a nonzero rate of 33%, is used as ground truth. Similar to before, we randomly masked values in the ground truth to generate the input, such that the resulting nonzero rate is 10%. We refer to the masked data set as PBMC_G949_10K (see summary statistics in **Table S1**).

### Data sets without the ground truth

We assessed the method performance also on the following real scRNA-seq data sets without the ground truth (**Table S1**; also see Data Availability):

1. A medium-sized PBMC data set (PBMC_G5561) of 5,561 genes in 53,970 cells, such that the total read count is >1K per gene and >1K per cell, and that the nonzero rate is 10%.
2. A large PBMC data set (PBMC_G9987) of 9,987 genes and 53,970 cells with the total read count >200 per gene and >1K per cell and a nonzero rate of 6%.
3. A smaller mouse brain data set from 10x Genomics (mouse_brain_G10K) of 10K genes and all the 1.3M cells with a nonzero rate of 20%.
4. The entire mouse brain data set (mouse_brain_G28K) of all the 27,998 genes and 1.3M cells with a nonzero rate of 7%.

### Principal Component Analysis (PCA)-based tSNE plots for visualization

When investigating the separation of cell types, we generated PCA-based tSNE plots to visualize how cells are grouped. To generate such plots, we first applied PCA to the data matrix, extracted the top principal components (e.g., 50 or 100), and then generated a tSNE plot using these top principal components (PCs) as the input. Since important features are summarized in top PCs, a tSNE plot based on these PCs provides a better visualization of key features than that based on the data matrix directly.

### Metrics for cell type separation

We calculate two metrics to assess the separation of cell types in the tNSE plots generated above: the between-cluster sum of squares (BCSS) and within-cluster sum of squares (WCSS). Denote the coordinates in the tSNE plot for the *i* th cell of the *k*th cell type (i.e., cluster) by **z**_*ki*_, which has length two, one for each dimension in the tSNE plot. Denote the number of cell types by *S*. Then we have the cluster mean as follows:

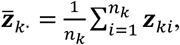

and the overall mean as follows:

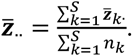

The sums of squares are defined as

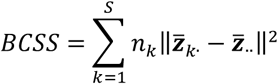

and

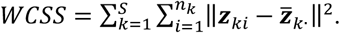

In the analysis of variance, BCSS is the between-treatment sum of squares, representing the source of variation due to different treatment groups (which are cell types here), whereas WCSS is the within-treatment sum of squares, representing the source of variation within a group. The sum of these two variations is the total sum of squares (TSS):

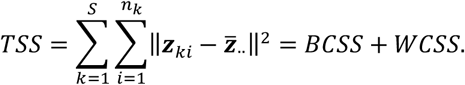

For the tSNE plot of cells here, BCSS measures the separation of cell types, and WCSS measures the closeness of cells within a cell type. In an extreme example, if all the cells are stacked on top of one another, then WCSS is zero, indicating extreme closeness within any cell type, and BCSS is also zero, indicating no separation of cell types. Conversely, clear cell type separation is indicated by large BCSS and small WCSS.

### Simulation with the R package Splatter

We used the R package Splatter (v1.6.1) to simulate scRNA-seq count data for 1000 genes and 20,000 cells from six clusters. The number of cells in each cluster ranges from nearly 700 to just above 6,000. We used the R function splatSimulate, letting the log normal distribution for the library size to have mean 15 and scale 0.2. For other parameters, we used the default values of this function. This step generated the ground truth. We then randomly masked 90% of the values with zero to create the input.

## ACKNOWLEDGEMENTS

We acknowledge the extensive support on hardware and software provided by the Computational Resources Core (CRC) through the Institute for Bioinformatics & Evolutionary Studies at the University of Idaho, and in particular the assistance from Dr. Benjamin Oswald, the Director of the CRC. This research is supported by NIH R00HG007368 (to A.Q.F.) and partially by the NIH/NIGMS grant P20GM104420 to the Center for Modeling Complex Interactions at the University of Idaho.

The Genotype-Tissue Expression (GTEx) Project was supported by the Common Fund of the Office of the Director of the National Institutes of Health, and by NCI, NHGRI, NHLBI, NIDA, NIMH, and NINDS. The gene expression data used for the analyses described in this manuscript were obtained from the GTEx Portal on 04/11/2018.

## ABBREVIATIONS

Adam: Adaptive Moment estimation
ALRA: Adaptively-thresholded Low-Rank Approximation
BCSS: Between-Cluster Sum of Squares
CPU: Central Processing Unit
DCA: Deep Count Autoencoder
GB: Gigabyte
GPU: Graphics Processing Unit
GTEx: Genotype-Tissue Expression
gtMSE: Mean Squared Error comparing with the ground truth
gtMSE_all_: Mean Squared Error comparing with the ground truth on all values
gtMSE_nz_: Mean Squared Error comparing with the ground truth only on nonzero values
LATE: Learning with AuToEncoder
MAGIC: Markov Affinity-based Graph Imputation of Cells
MSE: Mean Squared Error
PBMC: Peripheral Blood Mononuclear Cell
PC: Principal Component
PCA: Principal Component Analysis
RAM: Random Access Memory
ReLU: Rectified Linear Unit
SAVER: Single-cell Analysis Via Expression Recovery
scRNA-seq: single-cell RNA-sequencing
scVI: single-cell Variational Inference
SVD: Singular Value Decomposition
TB: Terabyte
TRANSLATE: TRANSfer learning with LATE
tSNE: t-distributed Stochastic Neighbor Embedding
TSS: Total Sum of Squares
WCSS: Within-Cluster Sum of Squares

## AUTHOR CONTRIBUTIONS

A.Q.F. conceived the project. R.L., B.L. and A.Q.F. developed the method. R.L., M.X. and A.Q.F. implemented the software. M.B.B., R.L. and A.Q.F. analyzed the data. M.B.B., R.L., B.L., Y.I.L., N.E.B. and A.Q.F. interpreted the results. A.Q.F. wrote the manuscript with input from other authors.

## COMPLIANCE WITH ETHICS GUIDELINES

The authors Md. Bahadur Badsha, Rui Li, Boxiang Liu, Yang I. Li, Min Xian, Nicholas E. Banovich and Audrey Qiuyan Fu declare that they have no conflicts of interest.

## DATA AVAILABILITY

Data sets used in this paper are published and publicly available at the following URLs:

- Mouse bone marrow data: GEO GSE72857. https://www.ncbi.nlm.nih.gov/geo/query/acc.cgi?acc=GSE72857.
- GTEx gene expression data (Gene TPMs): https://gtexportal.org/home/datasets.
- Human PBMC and mouse brain data from 10x Genomics: https://support.10xgenomics.com/single-cell-gene-expression/datasets.
- Mouse retina data: provided in the scVI software package (https://github.com/YosefLab/scVI/tree/master/tests/data); original data from GEO GSE81905. https://www.ncbi.nlm.nih.gov/geo/query/acc.cgi?acc=GSE81905.

## CODE AVAILABILITY

Our software is implemented in Python and builds on Google TensorFlow. It can be run on CPUs and GPUs. The source code is available at https://github.com/audreyqyfu/LATE.

