## Supplemental Figures for "Imputation of single-cell gene expression with an autoencoder neural network"

**Supplementary Figures**

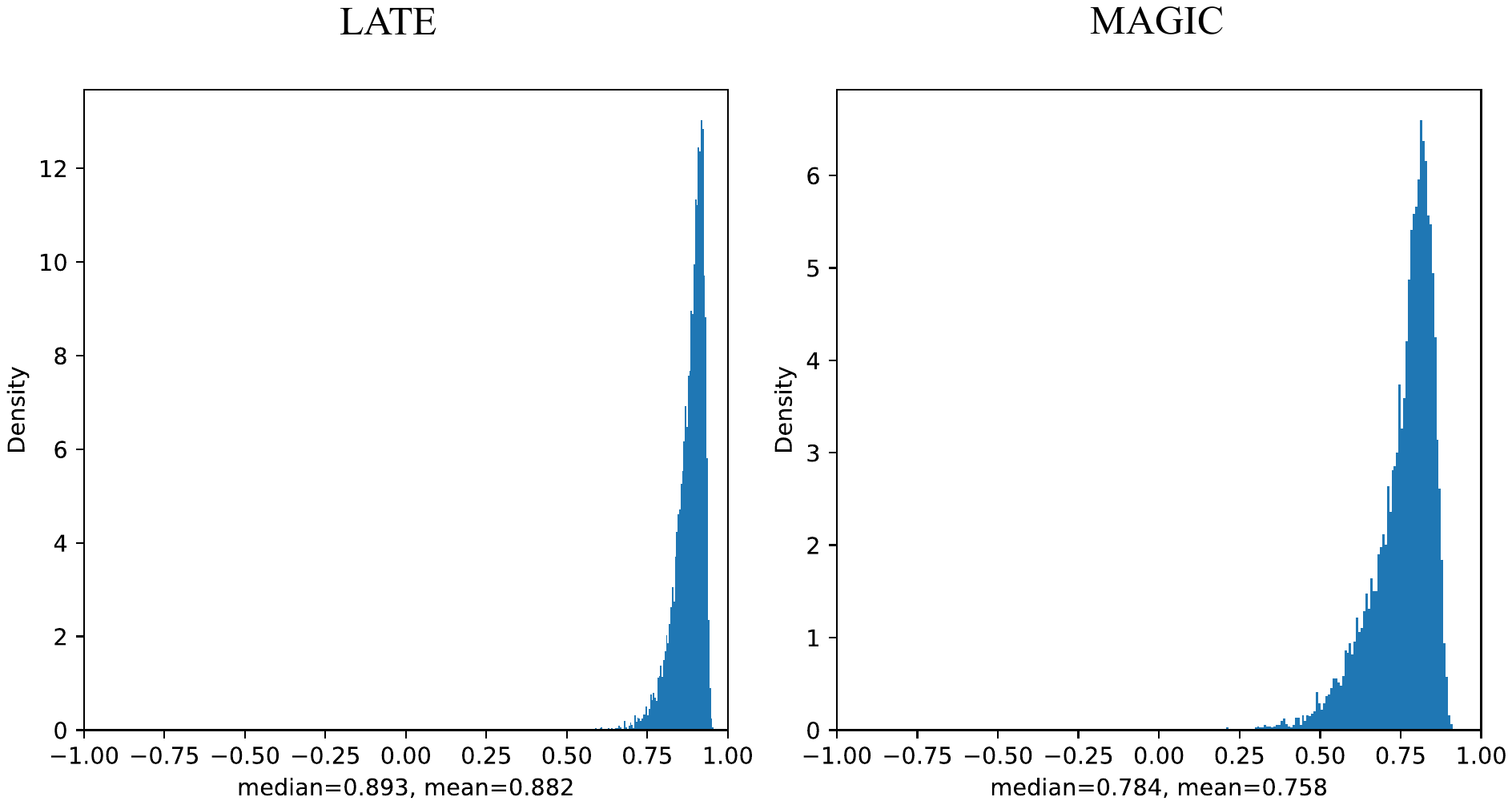

**Figure S1**: Histograms of Pearson correlations between the imputed gene expression profile and the ground truth in individual cells on the PBMC_G949_10K data (949 genes and 10K cells).

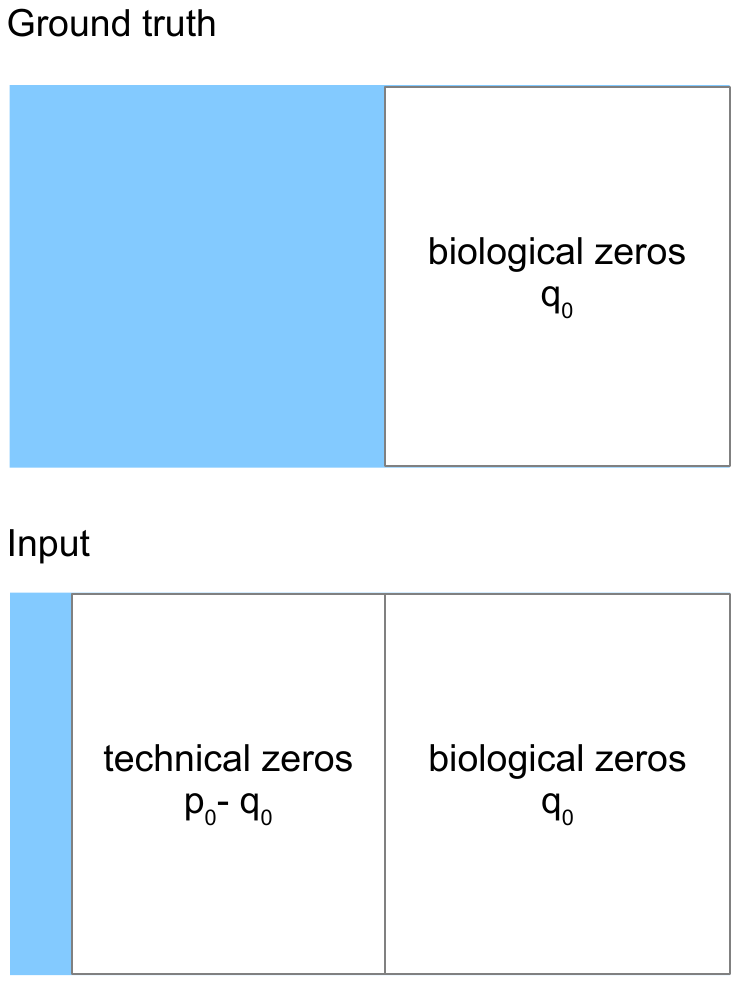

**Figure S2**: An illustration of biological and technical zeros in the ground truth and input. $p_{0}$ is the percentage of zeros in the input, and $q_{0}$ that in the ground truth.

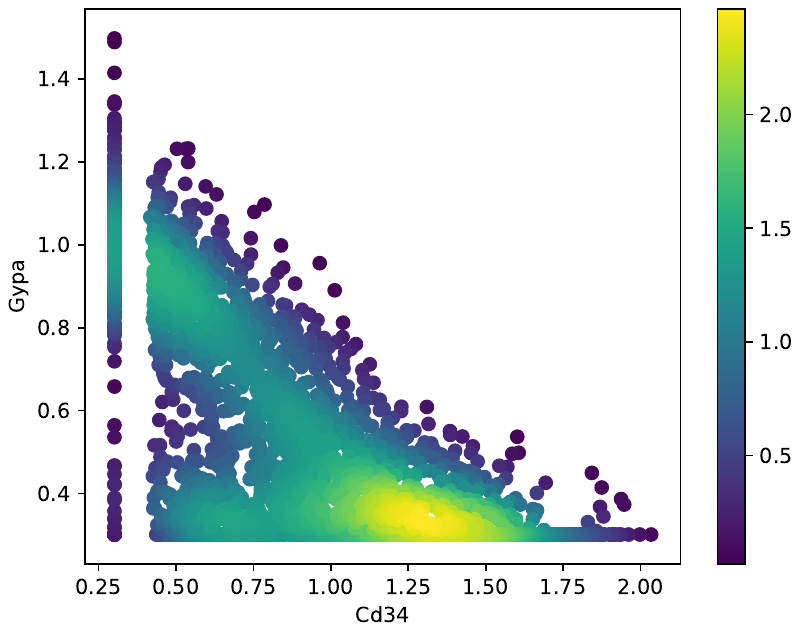

**Figure S3**: Two genes (*Cd34* vs *Gypa*) with a nonlinear relationship in the synthetic data based on the mouse bone marrow data (MAGIC_mouse; 16,114 genes and 2,576 cells) imputed by ALRA. Each dot is a single cell. The color bar indicates the Gaussian kernel density estimates of data points.

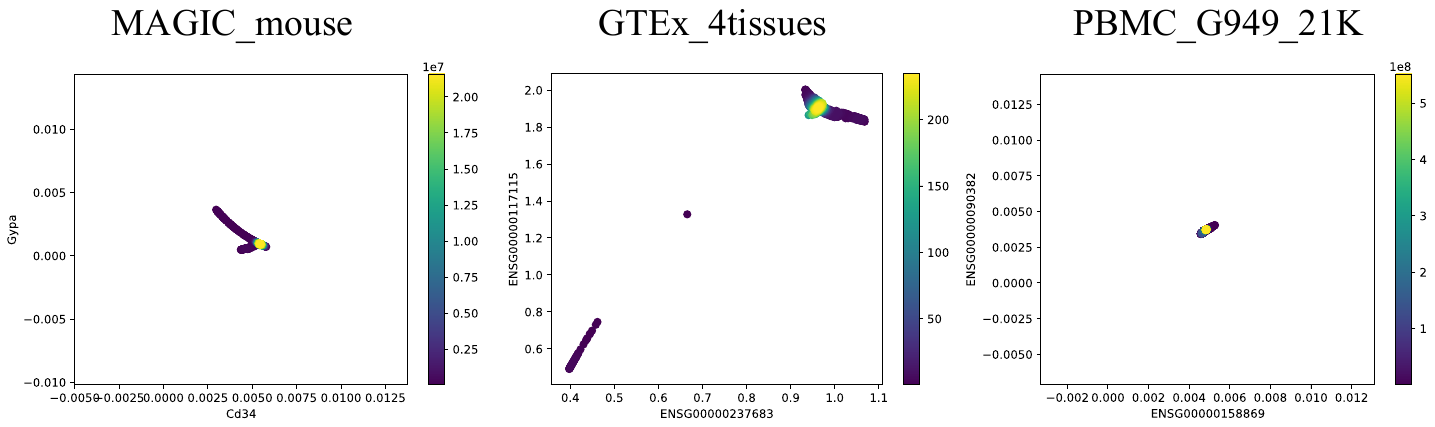

**Figure S4**: Nonlinear relationship results from MAGIC on the mouse bone marrow data (MAGIC_mouse; 16,114 genes and 2,576 cells), GTEx (GTEx_4tissues; 56,202 genes and 3,164 samples), and 10x Genomics PBMC data (PBMC_G949; 949 genes and 21,065 cells). Each dot is a single cell. The color bar indicates the Gaussian kernel density estimates of data points.

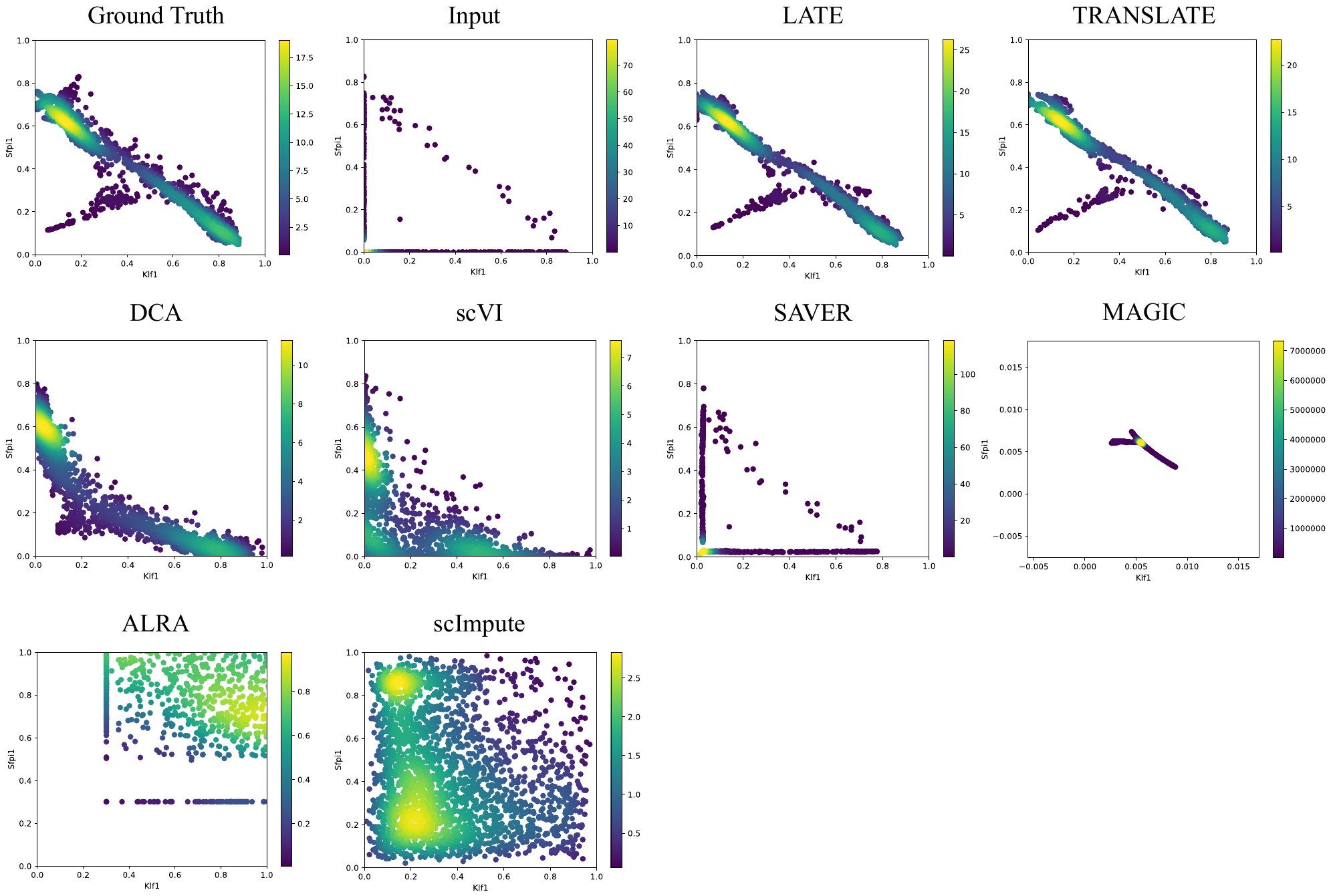

**Figure S5**: Two genes (*Klf1* vs *Sfpi1*) with a nonlinear relationship in the synthetic data based on the mouse bone marrow data (MAGIC_mouse; 16,114 genes and 2,576 cells). Each dot is a single cell. The color bar indicates the Gaussian kernel density estimates of data points.

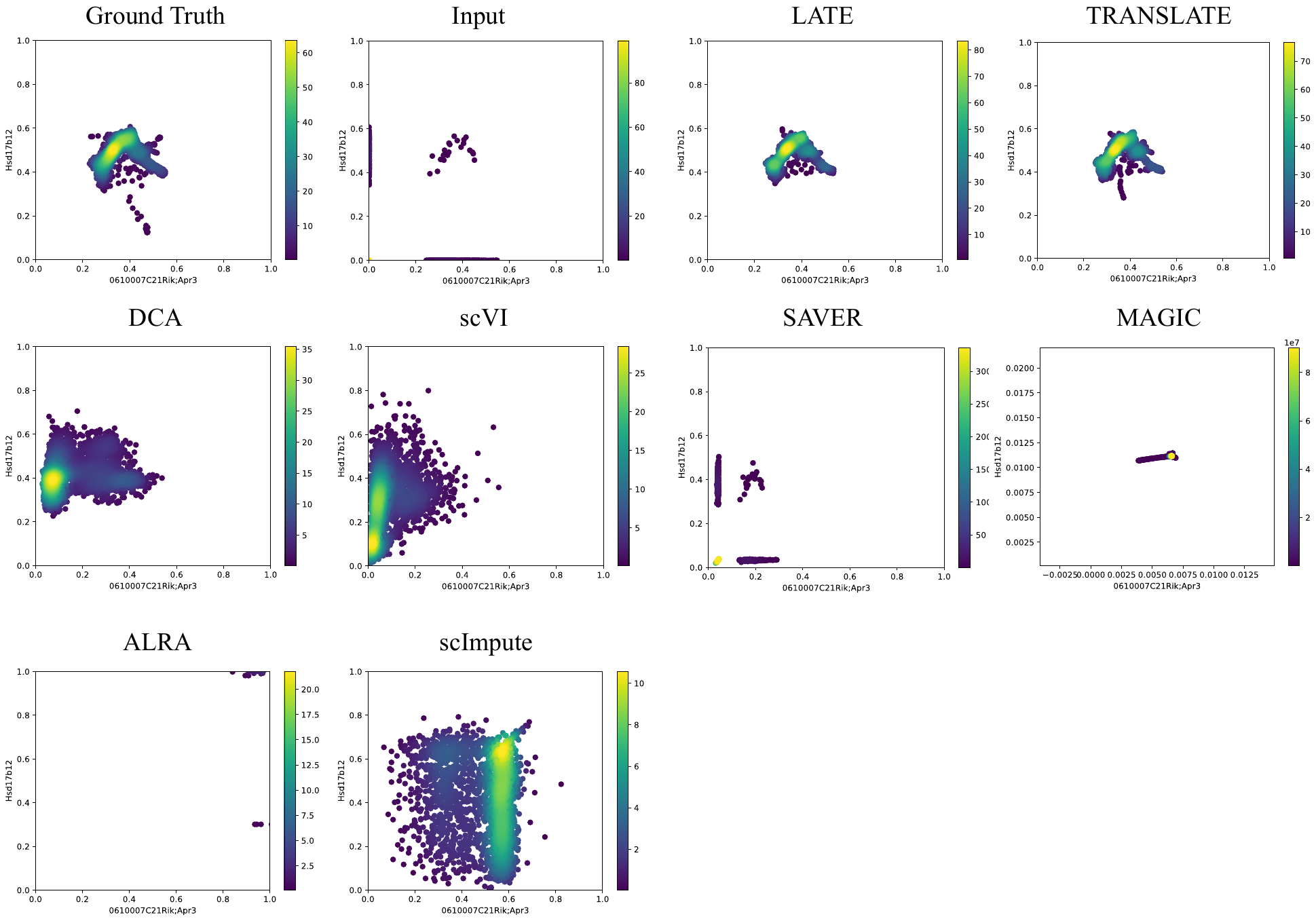

**Figure S6**: Two genes (*0610007C21Rik; Apr3* vs *Hsd17b12*) with a nonlinear relationship in the synthetic data based on the mouse bone marrow data (MAGIC_mouse; 16,114 genes and 2,576 cells). Each dot is a single cell. The color bar indicates the Gaussian kernel density estimates of data points.

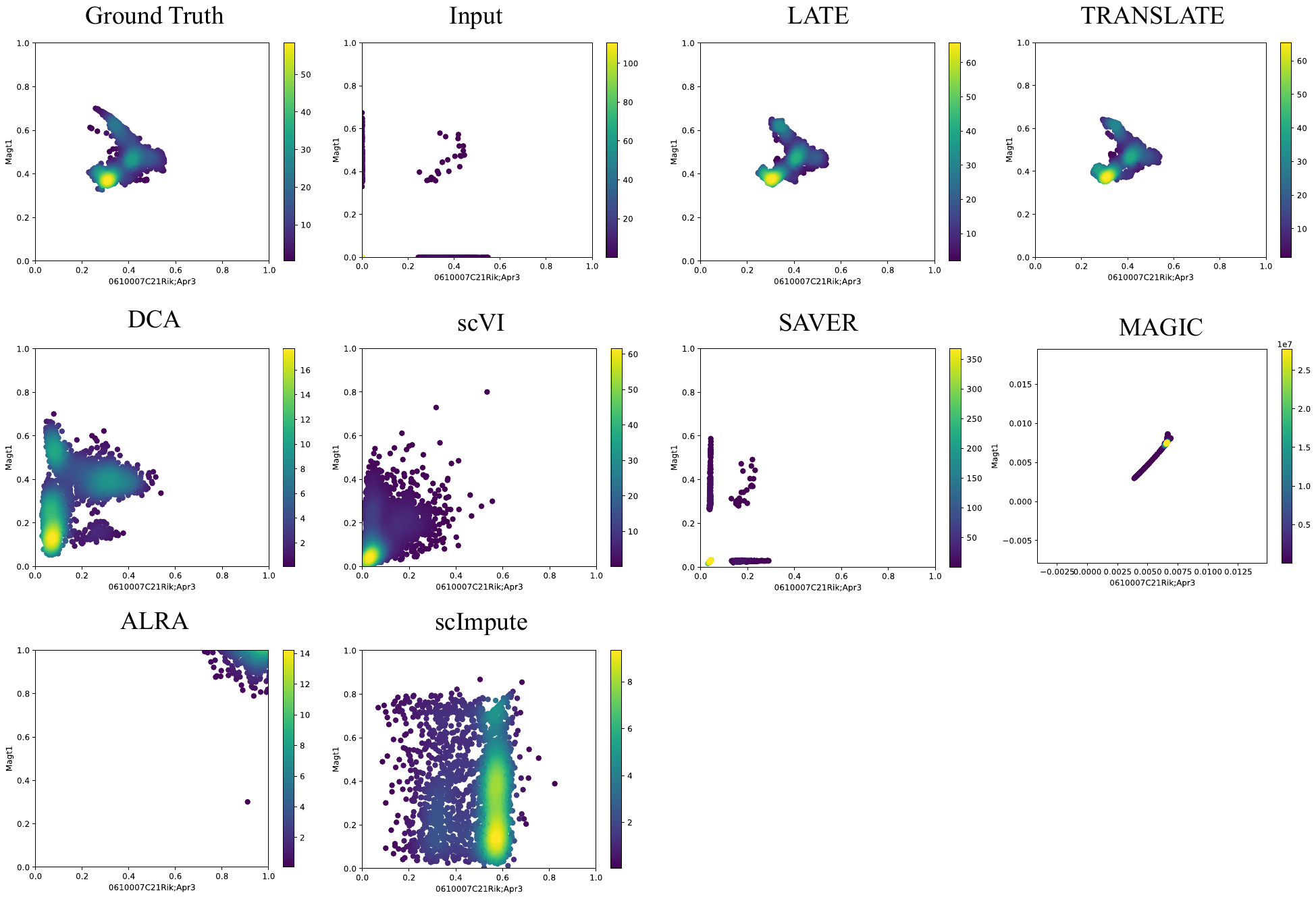

**Figure S7**: Two genes (*0610007C21Rik; Apr3* vs *Magt1*) with a nonlinear relationship in the synthetic data based on the mouse bone marrow data (MAGIC_mouse; 16,114 genes and 2,576 cells). Each dot is a single cell. The color bar indicates the Gaussian kernel density estimates of data points.

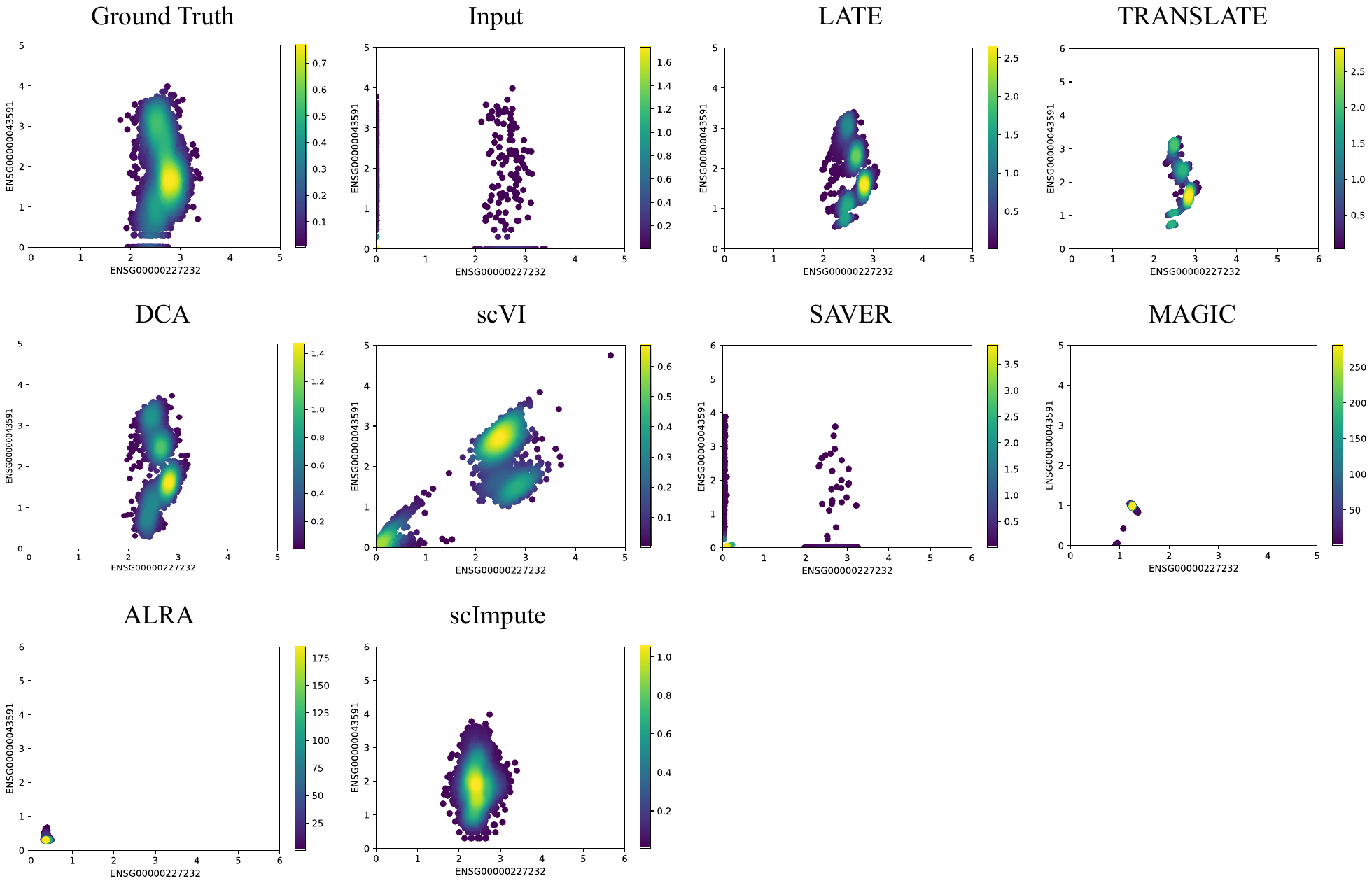

**Figure S8**: Two genes (*ENSG00000227232* vs *ENSG00000043591*) with a nonlinear relationship in the synthetic data based on GTEx (GTEx_4tissues; 56,202 genes and 3,164 samples). Each dot is a single cell. The color bar indicates the Gaussian kernel density estimates of data points.

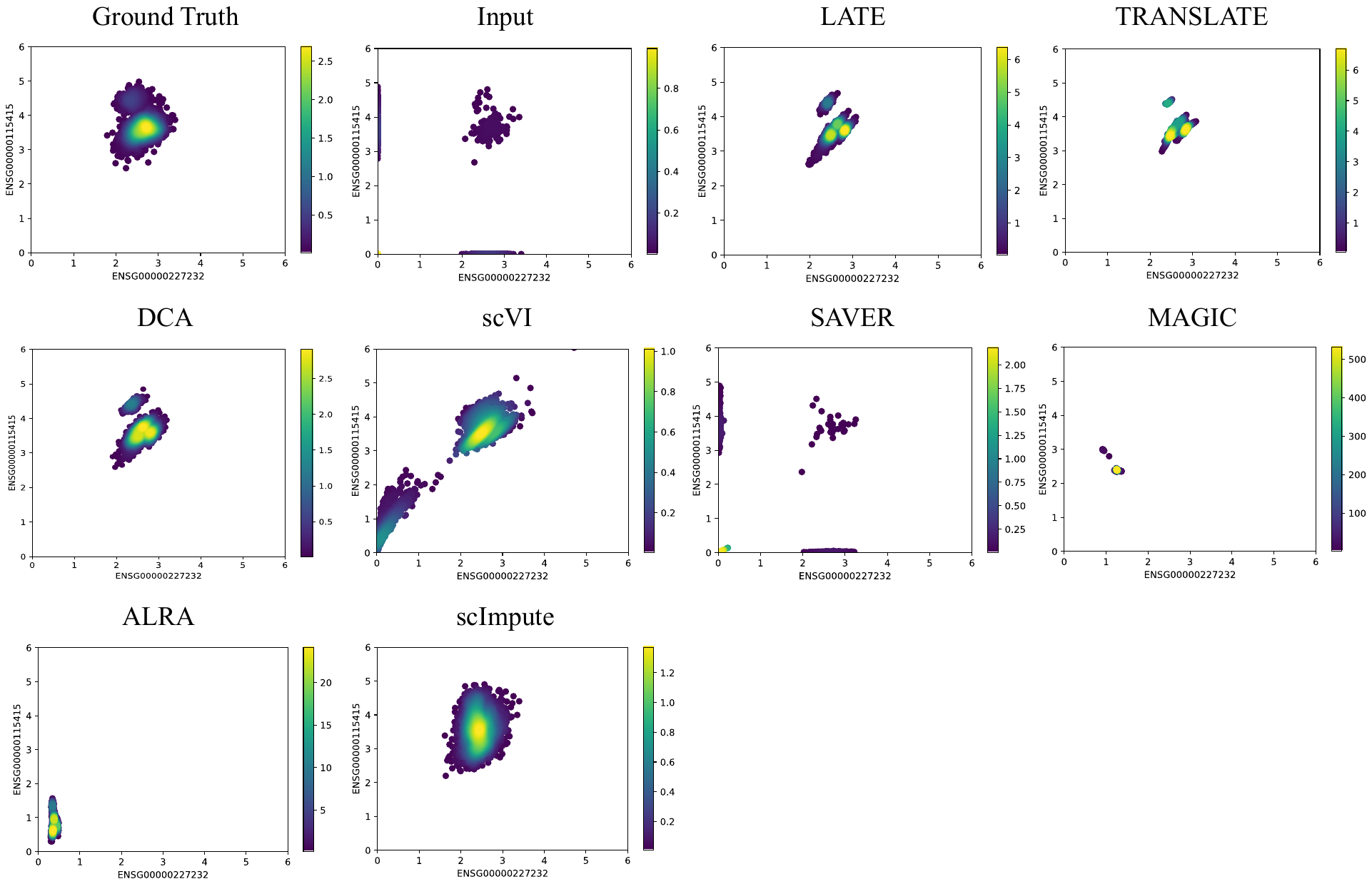

**Figure S9**: Two genes (*ENSG00000227232* vs *ENSG00000115415*) with a nonlinear relationship in the synthetic data based on GTEx (GTEx_4tissues; 56,202 genes and 3,164 samples). Each dot is a single cell. The color bar indicates the Gaussian kernel density estimates of data points.

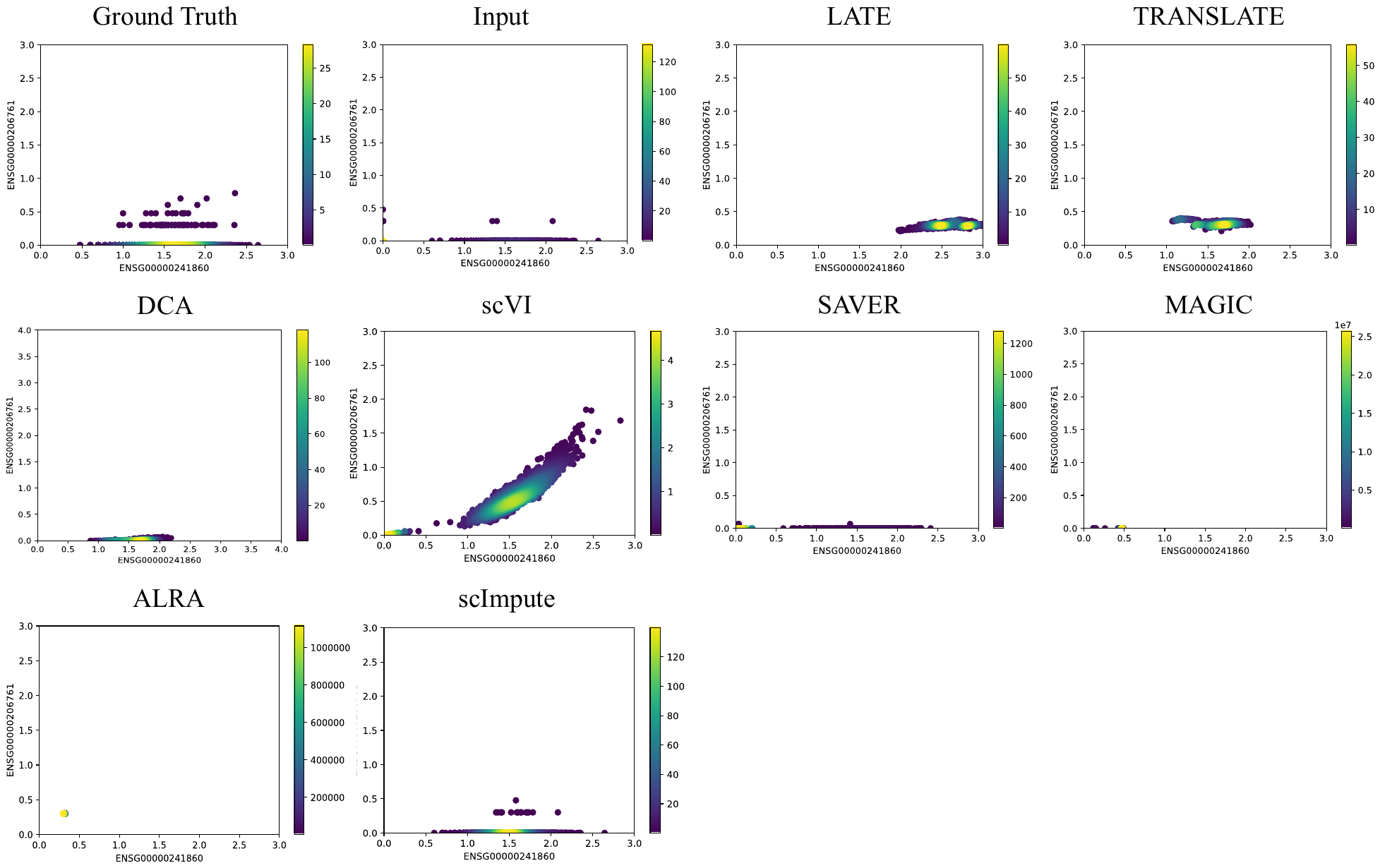

**Figure S10**: Two genes (*ENSG00000241860* vs *ENSG00000206761*) with a nonlinear relationship in the synthetic data based on GTEx (GTEx_4tissues; 56,202 genes and 3,164 samples). Each dot is a single cell. The color bar indicates the Gaussian kernel density estimates of data points.

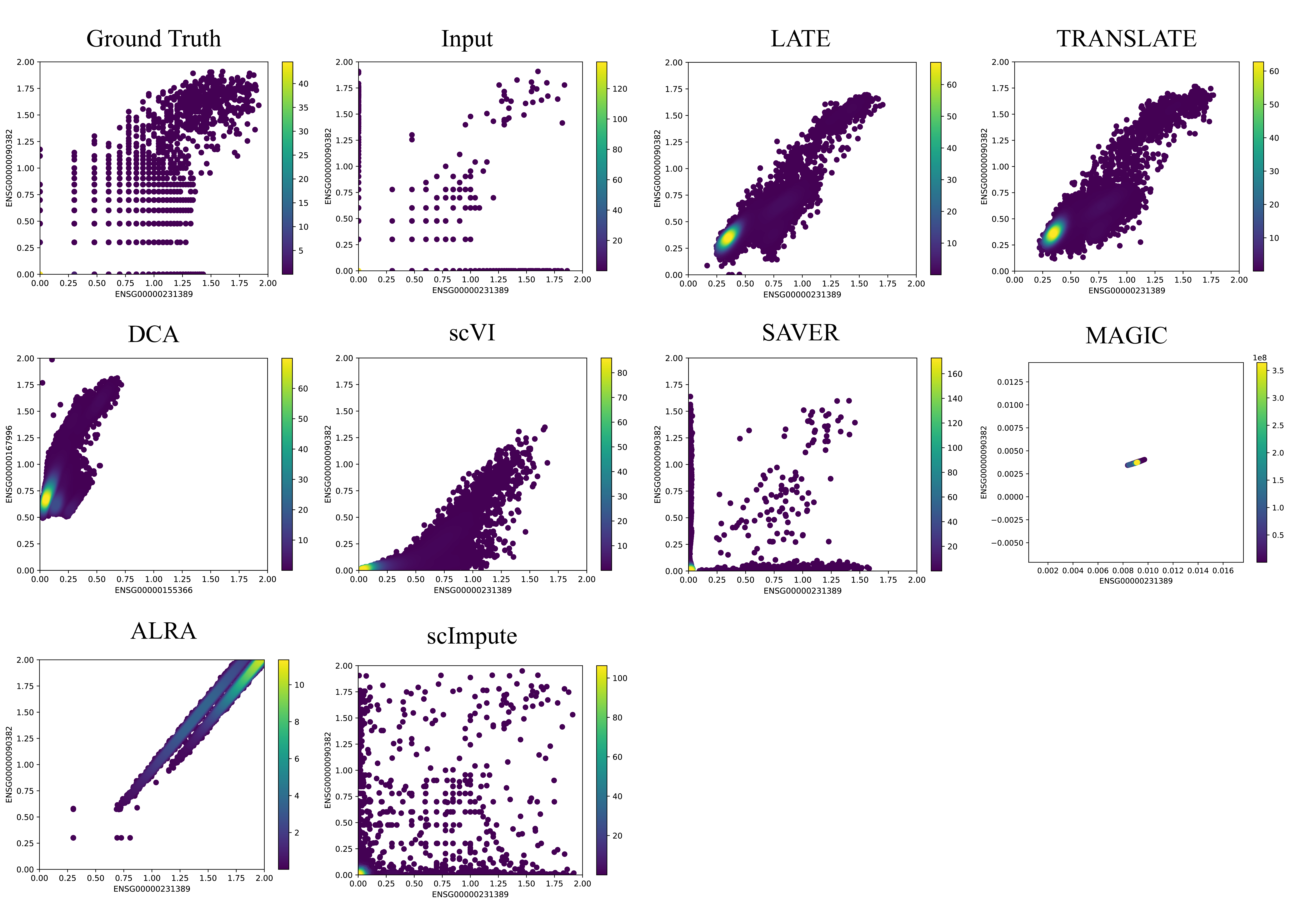

**Figure S11**: Two genes (*ENSG00000231389* vs *ENSG00000090382*) with a nonlinear relationship in the small synthetic data based on the 10x Genomics PBMC data (PBMC_G949; 949 genes and 21,065 cells). Each dot is a single cell. The color bar indicates the Gaussian kernel density estimates of data points.

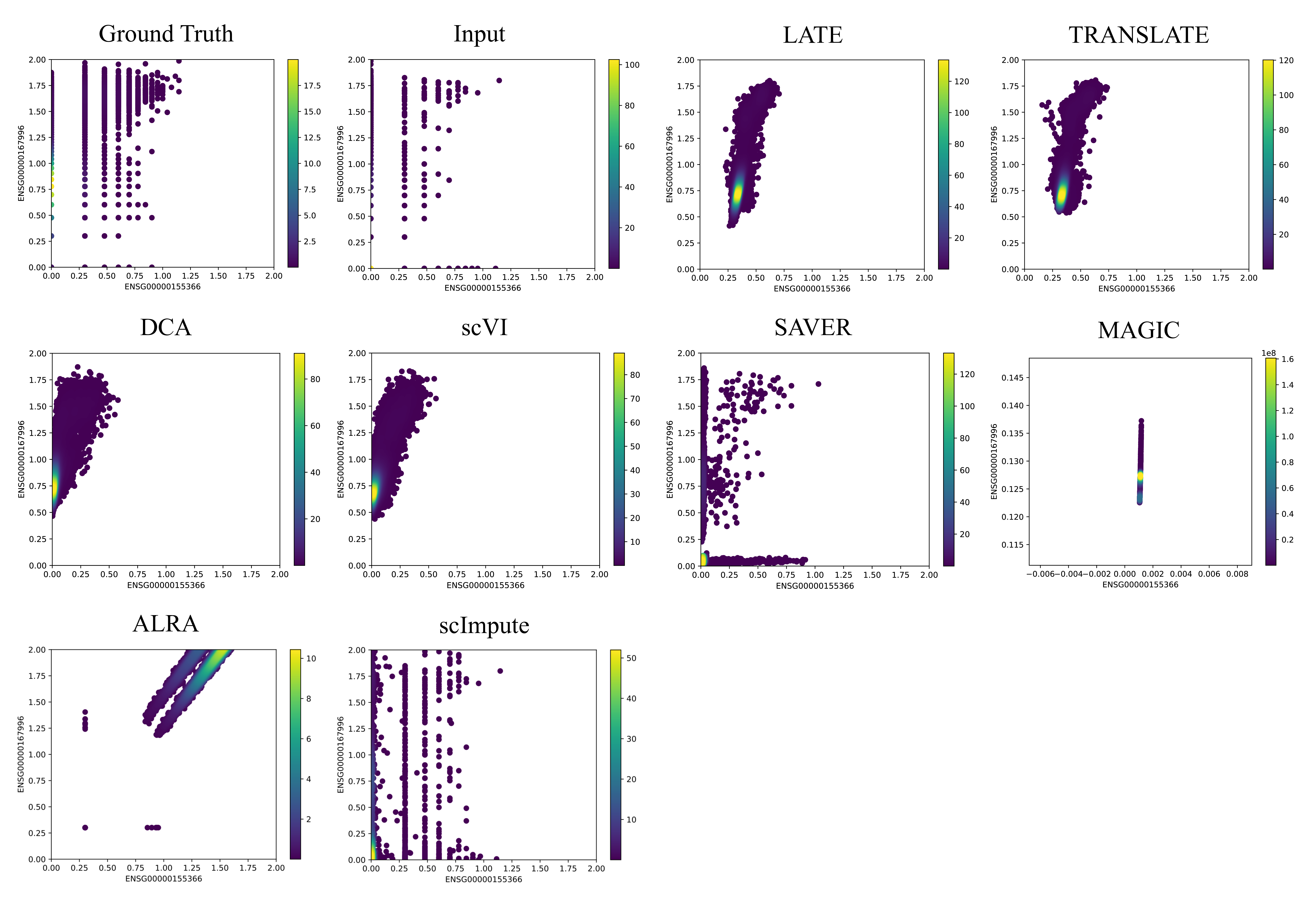

**Figure S12**: Two genes (*ENSG00000155366* vs *ENSG00000167996*) with a nonlinear relationship based on the 10x Genomics PBMC data (PBMC_G949; 949 genes and 21,065 cells). Each dot is a single cell. The color bar indicates the Gaussian kernel density estimates of data points.

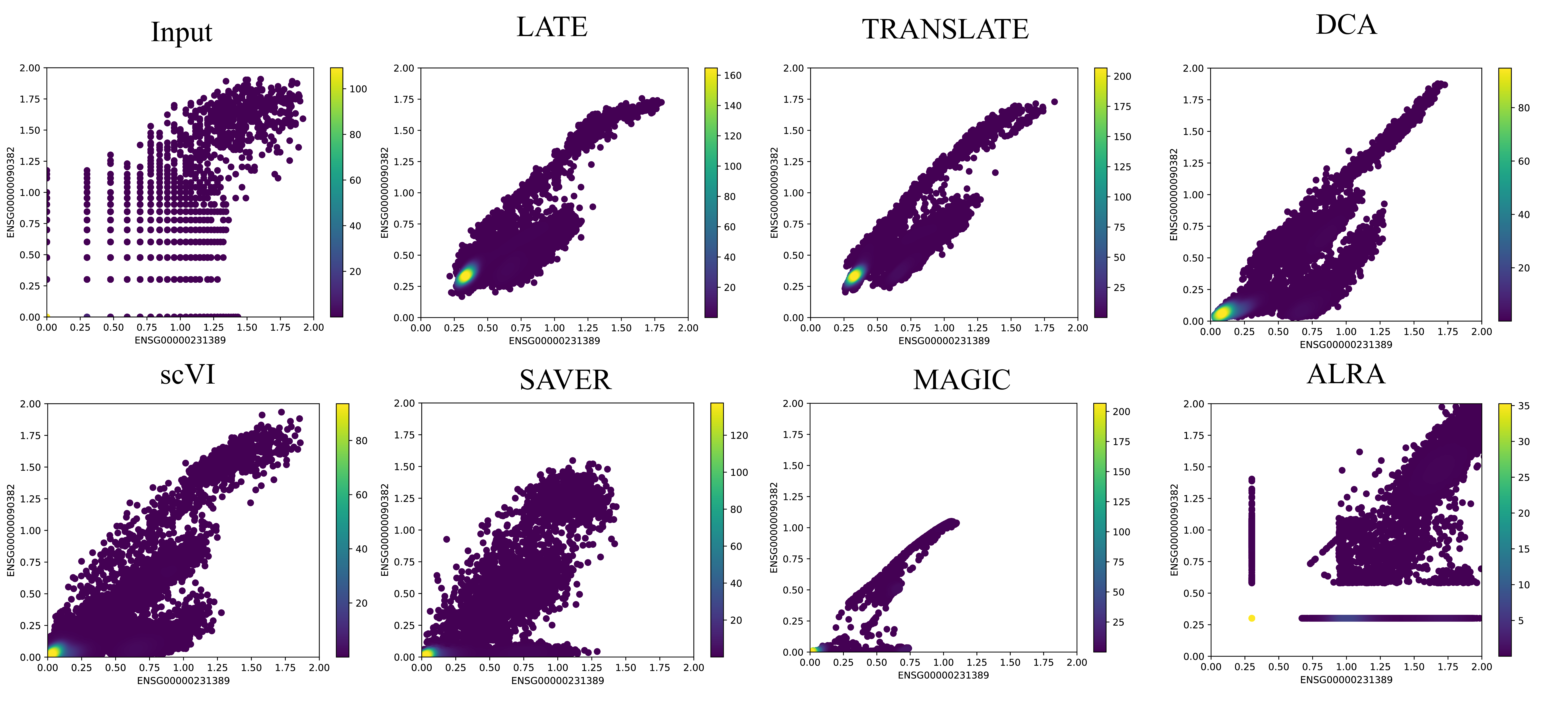

**Figure S13**: Two genes (*ENSG00000231389* vs *ENSG00000090382*) with a nonlinear relationship based on the 10x Genomics PBMC data (PBMC_G5561; 5,561 genes and 53,970 cells). Each dot is a single cell. The color bar indicates the Gaussian kernel density estimates of data points.

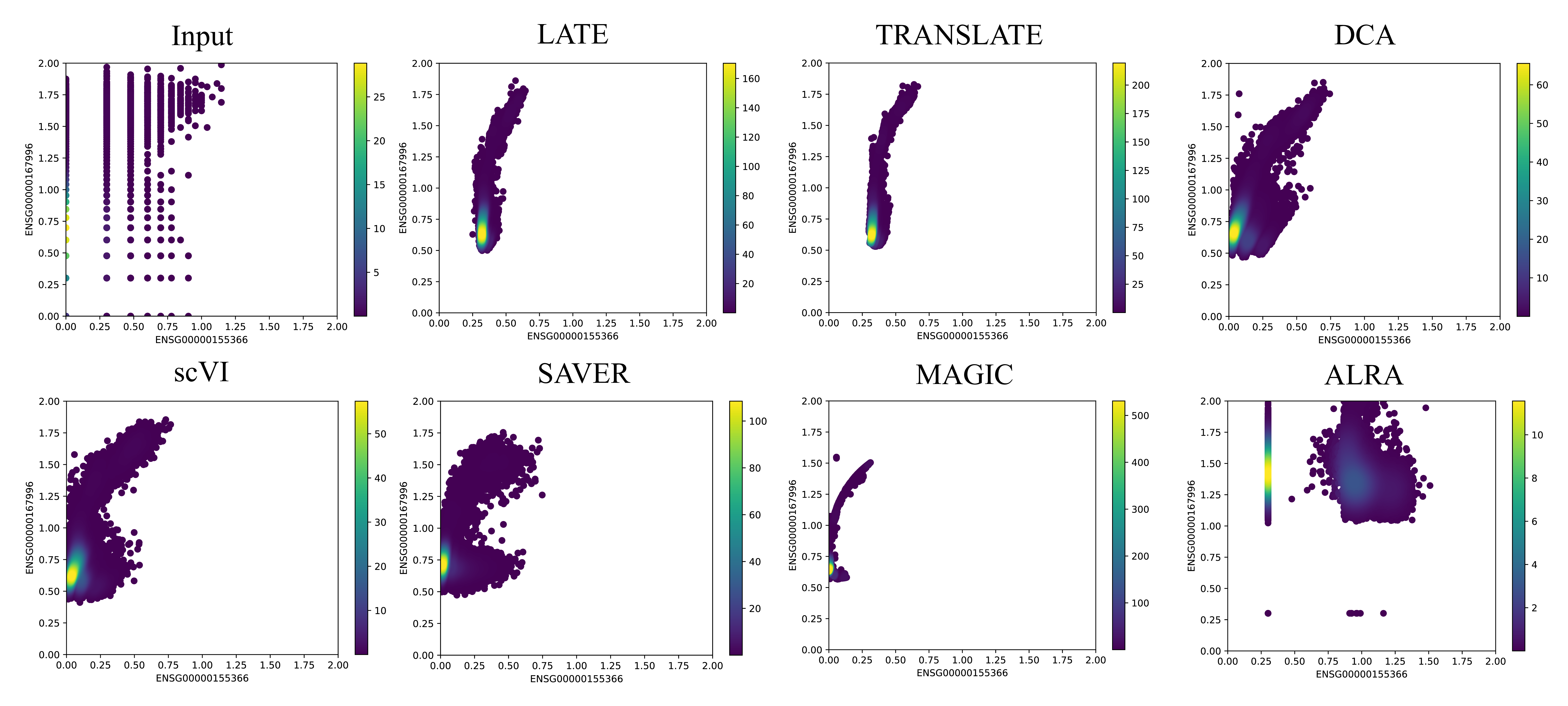

**Figure S14**: Two genes (*ENSG00000155366* vs *ENSG00000167996*) with a nonlinear relationship based on the 10x Genomics PBMC data (PBMC_G5561; 949 genes and 53,970 cells). Each dot is a single cell. The color bar indicates the Gaussian kernel density estimates of data points.

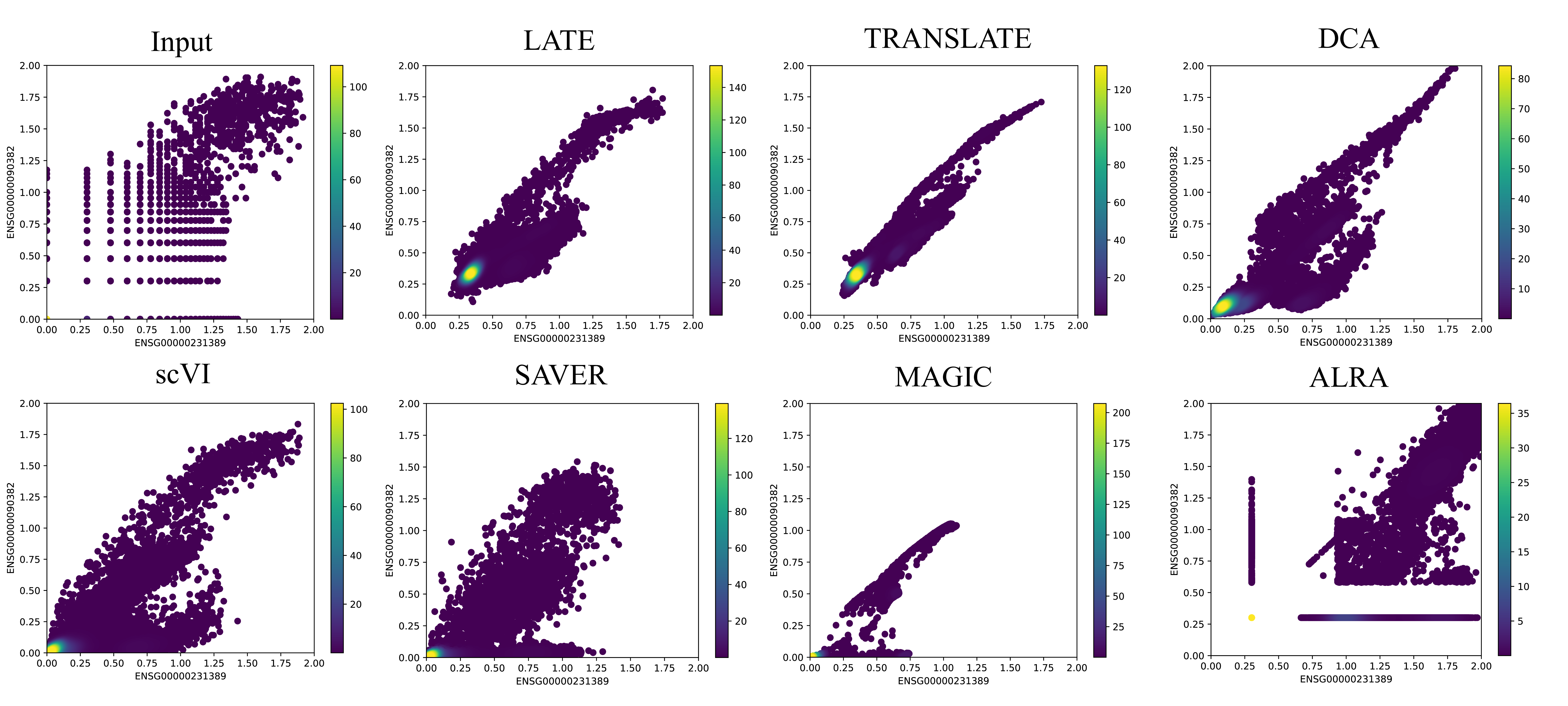

**Figure S15**: Two genes (*ENSG00000231389* vs *ENSG00000090382*) with a nonlinear relationship based on the 10x Genomics PBMC data (PBMC_G9987; 949 genes and 53,970 cells). Each dot is a single cell. The color bar indicates the Gaussian kernel density estimates of data points.

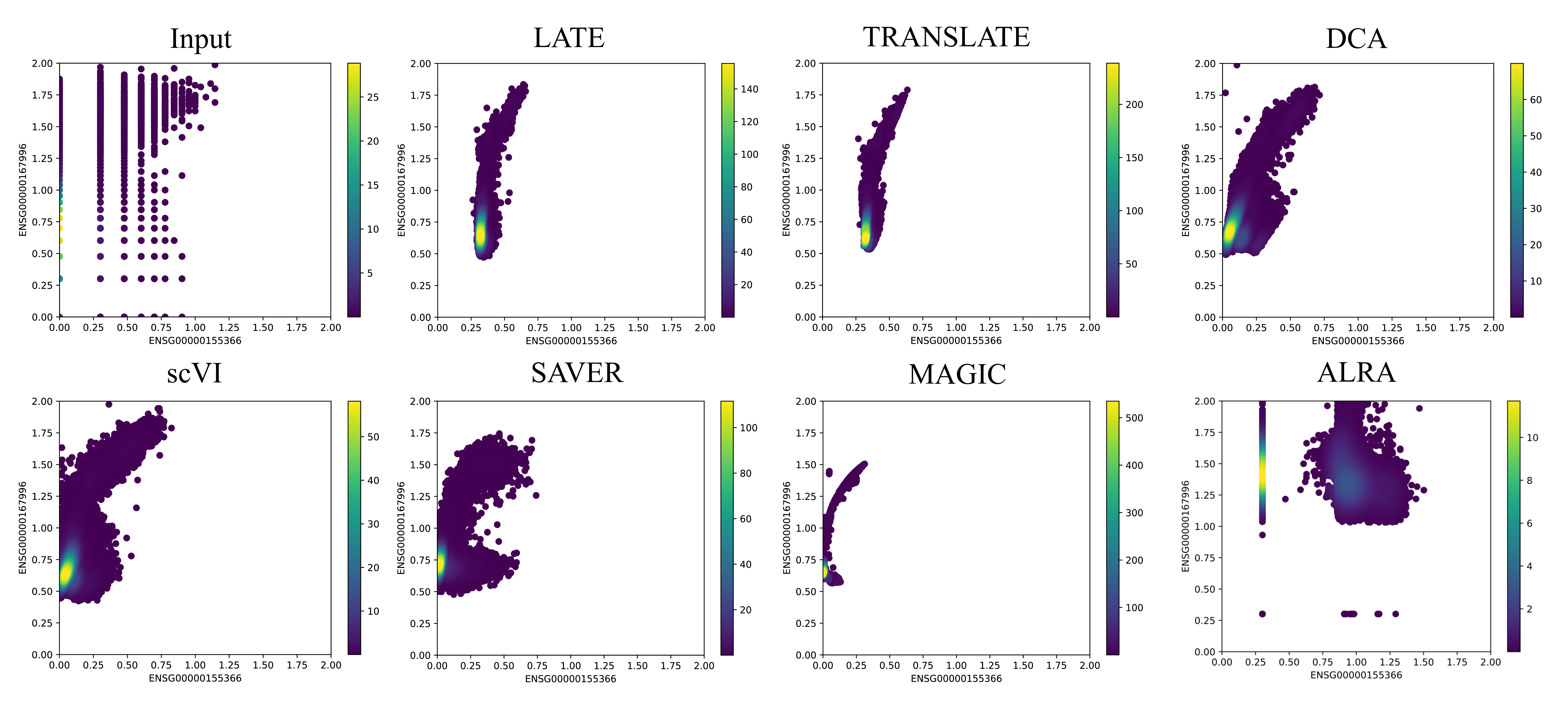

**Figure S16**: Two genes (*ENSG00000155366* vs *ENSG00000167996*) with a nonlinear relationship based on the 10x Genomics PBMC data (PBMC_G9987; 949 genes and 53,970 cells). Each dot is a single cell. The color bar indicates the Gaussian kernel density estimates of data points.

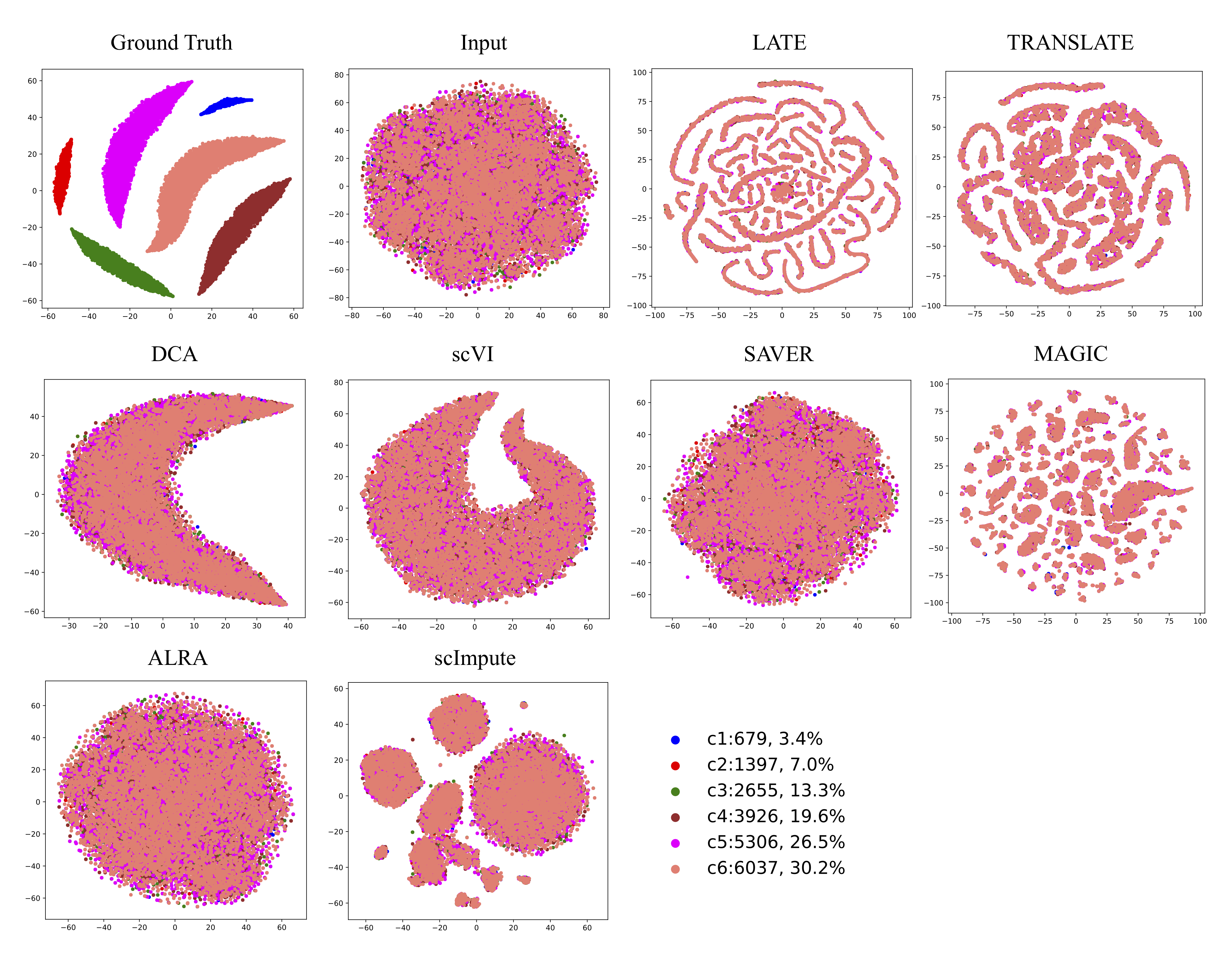

**Figure S17**. tSNE plots of cells from the synthetic data generated using the R package Splatter (1,000 genes, 20,000 cells, and 6 cell types). Each dot is a single cell, and each color indicates a cell type.

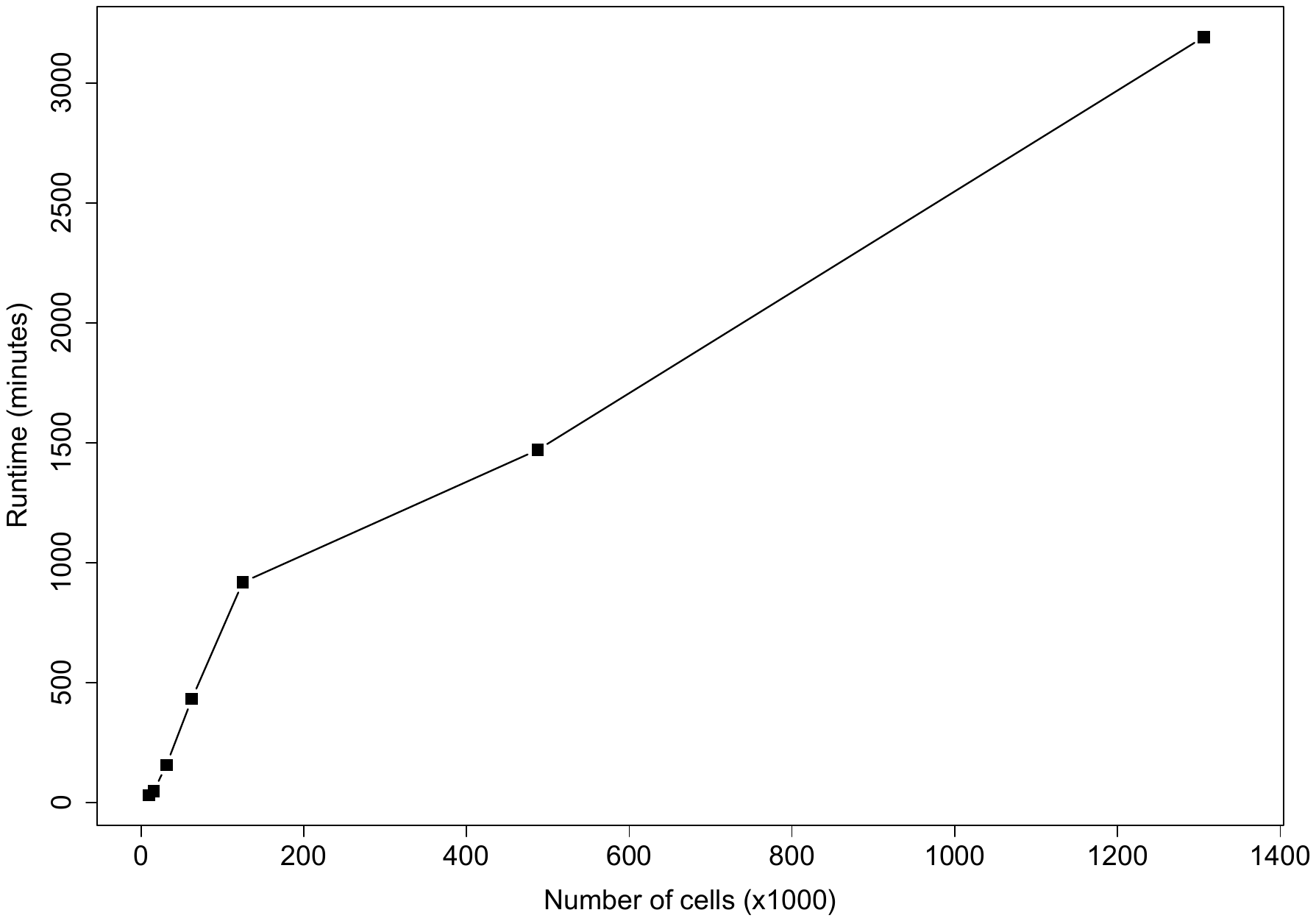

**Figure S18:** CPU runtime of LATE (genes as features) on multiple subsets of the mouse brain scRNA-seq data from 10x Genomics. The entire data set contains 28K genes and 1.3M cells. We sampled a subset of 10K genes with multiple subsets of cells as the input.
