## Supplemental Tables for "Imputation of single-cell gene expression with an autoencoder neural network"

**Supplementary Table S1: Description and summary statistics of data sets analyzed in the main text**.

| Data Set | Gene Count | Cell/Sample Count | Nonzero Rate | Ground Truth | Nonzero Rate in Ground Truth |
|---|---|---|---|---|---|
| MAGIC_mouse | 16,114 | 2,576 | 10% | Y | 100% |
| GTEx_4tissues | 56,202 | 3,164 | 10% | Y | 49.80% |
| PBMC_G949 | 949 | 21,065 | 10% | Y | 40.90% |
| PBMC_G5561 | 5,561 | 53,970 | 9.90% | N | NA |
| PBMC_G9987 | 9,987 | 53,970 | 5.90% | N | NA |
| mouse_brain_G10K | 10,000 | 1,306,111 | 19.65% | N | NA |
| mouse_brain_G28K | 27,998 | 1,306,127 | 7.18% | N | NA |
| PBMC_G949_10K | 949 | 10,000 | 10% | Y | 33% |

**Supplementary Table S2: Description and summary statistics of reference data sets used for TRANSLATE.**

| Input | Reference | | | |
|---|---|---|---|---|
| | Data Set | Type | Sample Size | Nonzero Rate |
| MAGIC_mouse | Mouse data following MAGIC imputation | single-cell | 6,182 | 99.80% |
| GTEx_4tissues | i) GTEx data with all tissues | bulk | 11,688 | 51.30% |
| GTEx_4tissues | ii) GTEx data without the four tissues in input | bulk | 7,981 | 51.90% |
| PBMC_G949 | GTEx data with all tissues | bulk | 11,688 | 51.30% |
| PBMC_G5561 | GTEx data with all tissues | bulk | 11,688 | 51.30% |
| PBMC_G9987 | GTEx data with all tissues | bulk | 11,688 | 51.30% |
| PBMC_G949_10K | i) 30K PBMCs with cell type labels | single-cell | 30,000 | 33% |
| PBMC_G949_10K | ii) PBMC data from another sample collection of the same individual | single-cell | 20,202 | 38.80% |
| PBMC_G949_10K | iii) 30K PBMCs with cell type labels and masking | single-cell | 30,000 | 10% |
| PBMC_G949_10K | iv) GTEx data with all tissues | bulk | 11,688 | 51.30% |

**Supplementary Table S3: MSEs (×10$^{-3}$) from different methods on multiple data sets**. See **Supplementary Tables S1 and S2** for description of the input data sets, as well as the reference data sets used for TRANSLATE. MSE is between imputation and input, and gtMSE between imputation and ground truth. In generally, $gtMSE_{nz}$ is reported, except for GTEx_4tissues, where zeros in the input are considered as real expression values and not missing values and therefore $gtMSE_{all}$ is reported.

| | **LATE (genes as features)** | | **LATE (combined)** | | **TRANSLATE (genes as features)** | | **TRANSLATE (combined)** | |
|---|---|---|---|---|---|---|---|---|
| **Data Set** | **MSE** | **gtMSE** | **MSE** | **gtMSE** | **MSE** | **gtMSE** | **MSE** | **gtMSE** |
| MAGIC_mouse | 0.07 | 0.89 | 0.07 | 0.90 | 0.03 | 0.44 | 0.03 | 0.44 |
| GTEx_4tissues i) | 5.18 | 104.57 | 5.14 | 113.45 | 5.37 | 114.82 | 5.30 | 117.97 |
| GTEx_4tissues ii) | | | | | 5.76 | 115.26 | 5.66 | 118.95 |
| PBMC_G949 | 1.56 | 7.23 | 1.56 | 7.23 | 1.88 | 7.84 | 1.56 | 7.26 |
| PBMC_G5561 | 1.11 | NA | 1.11 | NA | 1.10 | NA | 1.10 | NA |
| PBMC_G9987 | 0.63 | NA | 0.63 | NA | 0.65 | NA | 0.62 | NA |
| mouse_brain_G10K | 8.20 | NA | Cannot run | | Not run | | Cannot run | |
| mouse_brain_G28K | 4.00 | NA | Cannot run | | Not run | | Cannot run | |
| PBMC_G949_10K i) | 1.50 | 5.91 | 1.50 | 5.95 | 1.44 | 5.87 | 1.43 | 5.89 |
| PBMC_G949_10K ii) | | | | | 1.43 | 5.76 | 1.42 | 5.80 |
| PBMC_G949_10K iii) | | | | | 1.43 | 5.88 | Not run | |
| PBMC_G949_10K iv) | | | | | 1.82 | 6.22 | Not run | |

| | **DCA (early Dec 2018)** | | **scVI (v0.2.2 Nov 2018)** | | **SAVER (v1.1.1) Oct 2018** | | **MAGIC (v1.4.0) Nov 2018** | |
|---|---|---|---|---|---|---|---|---|
| **Data Set** | **MSE** | **gtMSE** | **MSE** | **gtMSE** | **MSE** | **gtMSE** | **MSE** | **gtMSE** |
| MAGIC_mouse | 2.08 | 22.60 | 4.59 | 51.50 | 2.05 | 87.06 | 10.47 | 104.77 |
| GTEx_4tissues i) | 5.75 | 53.53 | 126.59 | 788.28 | 264.10 | 2357.73 | 130.14 | 650.19 |
| GTEx_4tissues ii) | | | | | | | | |
| PBMC_G949 | 4.75 | 21.60 | 7.32 | 31.70 | 2.16 | 107.42 | 30.57 | 125.92 |
| PBMC_G5561 | 0.46 | NA | 5.28 | NA | 4.38 | NA | 8.53 | NA |
| PBMC_G9987 | 2.81 | NA | 3.25 | NA | 2.68 | NA | 5.37 | NA |
| mouse_brain_G10K | 42 hours | | 8% training done after 56 hours | | Failed | | Failed | |
| mouse_brain_G28K | Not run | | Not run | | Failed | | Failed | |
| PBMC_G949_10K i) | 4.56 | 17.41 | 5.96 | 21.60 | 2.86 | 69.86 | 25.54 | 85.77 |
| PBMC_G949_10K ii) | | | | | | | | |
| PBMC_G949_10K iii) | | | | | | | | |
| PBMC_G949_10K iv) | | | | | | | | |

| | ALRA (Aug 2019) | | scImpute (Sep 2019) | |
|---|---|---|---|---|
| **Data Set** | **MSE** | **gtMSE** | **MSE** | **gtMSE** |
| MAGIC_mouse | 22.94 | 204.38 | 0.00 | 30.30 |
| GTEx_4tissues i) | 415.43 | 2097.14 | 0.30 | 137.00 |
| GTEx_4tissues ii) | | | | |
| PBMC_G949 | 163.30 | 652.93 | 0.00 | 73.70 |
| PBMC_G5561 | 50.45 | NA | Failed | |
| PBMC_G9987 | 28.84 | NA | Failed | |
| mouse_brain_G10K | Failed | | Not run | |
| mouse_brain_G28K | Failed | | Not run | |
| PBMC_G949_10K i) | 181.76 | 593.02 | 0.00 | 51.60 |
| PBMC_G949_10K ii) | | | | |
| PBMC_G949_10K iii) | | | | |
| PBMC_G949_10K iv) | | | | |

**Supplementary Table S4: Different types of gtMSEs ($\times 10^{-3}$) from all the methods on multiple data sets with ground truth**. See **Supplementary Tables S1 and S2** for description of the input data sets, as well as the reference data sets used for TRANSLATE. "all" refers to $gtMSE_{all}$, "nonzero" $gtMSE_{nz}$, "biol" $gtMSE_{biol}$, and "tech" $gtMSE_{tech}$. See "Assessing imputation accuracy" in Materials and Methods for definitions and calculations of the gtMSEs.

| | LATE (genes as features) | | | | LATE (combined) | | | |
|---|---|---|---|---|---|---|---|---|
| **Data Set** | **all** | **nonzero** | **biol** | **tech** | **all** | **nonzero** | **biol** | **tech** |
| MAGIC_mouse | 0.89 | 0.89 | NA | 0.98 | 0.90 | 0.90 | NA | 0.99 |
| GTEx_4tissues i) | 104.57 | 27.27 | 182.08 | 32.86 | 113.45 | 27.08 | 198.22 | 32.62 |
| GTEx_4tissues ii) | | | | | | | | |
| PBMC_G949 | 79.57 | 7.23 | 129.68 | 9.06 | 79.57 | 7.23 | 129.68 | 9.06 |
| PBMC_G949_10K i) | 81.95 | 5.91 | 119.48 | 7.83 | 82.00 | 5.95 | 119.46 | 7.88 |
| PBMC_G949_10K ii) | | | | | | | | |
| PBMC_G949_10K iii) | | | | | | | | |
| PBMC_G949_10K iv) | | | | | | | | |

| | TRANSLATE (genes as features) | | | | TRANSLATE (combined) | | | |
|---|---|---|---|---|---|---|---|---|
| **Data Set** | **all** | **nonzero** | **biol** | **tech** | **all** | **nonzero** | **biol** | **tech** |
| MAGIC_mouse | 0.44 | 0.44 | NA | 0.49 | 0.44 | 0.44 | NA | 0.49 |
| GTEx_4tissues i) | 114.82 | 28.02 | 201.31 | 33.69 | 117.97 | 27.69 | 207.58 | 33.33 |
| GTEx_4tissues ii) | 115.26 | 29.72 | 199.62 | 35.72 | 118.95 | 29.27 | 207.99 | 35.24 |
| PBMC_G949 | 86.36 | 7.84 | 140.77 | 9.77 | 78.89 | 7.26 | 128.48 | 9.10 |
| PBMC_G949_10K i) | 80.97 | 5.87 | 118.00 | 7.80 | 81.53 | 5.89 | 118.74 | 7.83 |
| PBMC_G949_10K ii) | 80.14 | 5.76 | 116.72 | 7.64 | 81.46 | 5.80 | 118.79 | 7.70 |
| PBMC_G949_10K iii) | 81.97 | 5.88 | 119.49 | 7.81 | | | | |
| PBMC_G949_10K iv) | 98.70 | 6.22 | 144.25 | 8.13 | | | | |

| | DCA | | | | scVI | | | |
|---|---|---|---|---|---|---|---|---|
| **Data Set** | **all** | **nonzero** | **biol** | **tech** | **all** | **nonzero** | **biol** | **tech** |
| MAGIC_mouse | 22.60 | 22.60 | NA | 24.88 | 51.50 | 51.50 | NA | 56.71 |
| GTEx_4tissues i) | 53.78 | 31.26 | 76.12 | 37.59 | 788.28 | 622.38 | 952.68 | 746.37 |
| GTEx_4tissues ii) | | | | | | | | |
| PBMC_G949 | 31.43 | 21.60 | 38.18 | 27.05 | 36.35 | 31.70 | 39.65 | 39.59 |
| PBMC_G949_10K i) | 26.21 | 17.41 | 30.53 | 22.98 | 27.62 | 21.60 | 30.56 | 28.40 |
| PBMC_G949_10K ii) | | | | | | | | |
| PBMC_G949_10K iii) | | | | | | | | |
| PBMC_G949_10K iv) | | | | | | | | |

| | SAVER | | | | MAGIC | | | |
|---|---|---|---|---|---|---|---|---|
| **Data Set** | **all** | **nonzero** | **biol** | **tech** | **all** | **nonzero** | **biol** | **tech** |
| MAGIC_mouse | 87.06 | 87.06 | NA | 96.55 | 104.77 | 104.77 | NA | 115.50 |
| GTEx_4tissues i) | 2357.75 | 2357.73 | 2360.00 | 2886.63 | 652.20 | 650.19 | 653.98 | 780.65 |
| GTEx_4tissues ii) | | | | | | | | |
| PBMC_G949 | 107.89 | 107.42 | 108.69 | 140.93 | 125.97 | 125.92 | 126.00 | 156.87 |
| PBMC_G949_10K i) | 70.54 | 69.86 | 70.80 | 99.05 | 85.88 | 85.77 | 85.95 | 112.02 |
| PBMC_G949_10K ii) | | | | | | | | |
| PBMC_G949_10K iii) | | | | | | | | |
| PBMC_G949_10K iv) | | | | | | | | |

<table>
<tr><th></th><th colspan="4">ALRA</th><th colspan="4">scImpute</th></tr>
<tr><th>Data Set</th><th>all</th><th>nonzero</th><th>biol</th><th>tech</th><th>all</th><th>nonzero</th><th>biol</th><th>tech</th></tr>
<tr><td>MAGIC_mouse</td><td>275.66</td><td>275.43</td><td>NA</td><td>302.14</td><td>30.28</td><td>30.28</td><td>NA</td><td>33.67</td></tr>
<tr><td>GTEx_4tissues i)</td><td rowspan="2">1811.90</td><td rowspan="2">1766.33</td><td rowspan="2">1849.69</td><td rowspan="2">2126.53</td><td rowspan="2">137.11</td><td rowspan="2">129.59</td><td rowspan="2">143.94</td><td rowspan="2">162.59</td></tr>
<tr><td>GTEx_4tissues ii)</td></tr>
<tr><td>PBMC_G949</td><td>2079.53</td><td>661.26</td><td>3062.01</td><td>821.19</td><td>75.77</td><td>73.74</td><td>77.25</td><td>97.55</td></tr>
<tr><td>PBMC_G949_10K i)</td><td rowspan="4">2276.13</td><td rowspan="4">599.24</td><td rowspan="4">3107.96</td><td rowspan="4">779.43</td><td rowspan="4">54.06</td><td rowspan="4">51.57</td><td rowspan="4">55.33</td><td rowspan="4">74.03</td></tr>
<tr><td>PBMC_G949_10K ii)</td></tr>
<tr><td>PBMC_G949_10K iii)</td></tr>
<tr><td>PBMC_G949_10K iv)</td></tr>
</table>

**Supplementary Table S5: GPU runtime of LATE (genes as features) on multiple subsets of the mouse brain scRNA-seq data from 10x Genomics.**

| Number of Genes ($\times 10^3$) | Number of Cells ($\times 10^3$) | GPU Runtime (Minutes) |
|---|---|---|
| 10 | 10.0 | 4.5 |
| 10 | 15.6 | 6.3 |
| 10 | 31.3 | 12.0 |
| 10 | 62.5 | 22.7 |
| 10 | 125.0 | 32.4 |
| 10 | 325.0 | 95.0 |
| 10 | 487.5 | 189.2 |
| 10 | 650.0 | 188.4 |
| 10 | 812.5 | 290.4 |
| 10 | 975.0 | 343.5 |
| 10 | 1306.1 | 440.2 |
| 28 | 10.2 | 10.7 |
| 28 | 15.6 | 14.9 |
| 28 | 40.6 | 57.2 |
| 28 | 162.5 | 94.1 |
| 28 | 325.0 | 197.9 |
| 28 | 487.5 | 289.4 |
| 28 | 650.0 | 382.1 |
| 28 | 812.5 | 476.3 |
| 28 | 975.0 | 553.0 |
| 28 | 1306.1 | 651.0 |

**Supplementary Table S6: CPU runtime of LATE (genes as features) on multiple subsets of the mouse brain scRNA-seq data from 10x Genomics.**

| Number of Genes ($\times 10^3$) | Number of Cells ($\times 10^3$) | CPU Runtime (Minutes) |
|---|---|---|
| 10 | 10.0 | 30 |
| 10 | 15.6 | 48 |
| 10 | 31.3 | 156 |
| 10 | 62.5 | 432 |
| 10 | 125.0 | 918 |
| 10 | 487.5 | 1470 |
| 10 | 1306.1 | 3192 |
